# Integrated single-cell transcriptomic and epigenetic analyses of cell-state transition and lineage commitment in the embryonic mouse cerebellum

**DOI:** 10.1101/2021.08.08.455565

**Authors:** Nagham Khouri-Farah, Qiuxia Guo, Kerry Morgan, Jihye Shin, James Y.H. Li

## Abstract

Recent studies using single-cell RNA-seq have revealed cellular heterogeneity in the developing mammalian cerebellum, yet the regulatory logic underlying this cellular diversity remains to be elucidated. Using integrated single-cell RNA and ATAC analyses, we resolved developmental trajectories of cerebellar progenitors and identified putative *trans*- and *cis*- elements that control cell state transition. We reverse-engineered gene regulatory networks (GRNs) of each cerebellar cell type. Through *in silico* simulations and *in vivo* experiments, we validated the efficacy of GRN analyses and uncovered the molecular control of a newly identified stem zone, the posterior transitory zone (PTZ), which contains multipotent progenitors for granule neurons, Bergmann glia, and choroid plexus epithelium. Importantly, we showed that perturbing cell fate specification of PTZ progenitors causes posterior cerebellar vermis hypoplasia, the most common cerebellar birth defect in humans. Our study provides a foundation for comprehensive studies of developmental programs of the mammalian cerebellum.

## INTRODUCTION

The cerebellum, which contains 80% of the total neurons of the human brain, is important in cognitive processing and sensory discrimination, in addition to its well-known function in motor coordination ^1^. There is a resurgence of interest in the development of the cerebellum as it is recognized as a locus for numerous developmental brain disorders, such as ataxia, autism, schizophrenia, and attention deficit-hyperactivity disorder ^2^.

The embryonic cerebellum contains two spatially distinct germinal zones: the ventricular zone (VZ) and the upper rhombic lip (RL), which give rise to inhibitory and excitatory neurons, respectively ^3^. From these germinal zones, different types of GABAergic and glutamatergic neurons are generated in temporally restricted phases ^3^. In particular, the RL produces cerebellar glutamatergic nuclear neurons between E9.5 and E12.5, granule cells after E12.5, and unipolar brush cells at E16.5 ^4, 5^. Genetic studies have demonstrated that transcription factor (TF) Ptf1a controls the GABAergic fate of VZ progenitors, whereas Atoh1 controls the glutamatergic fate of RL cells ^6–9^. Despite these advances, the gene regulatory programs in each cerebellar cell type remain to be defined.

It was suggested that RL-derived *Atoh1* cells are induced *de novo* throughout embryogenesis ^5^. Currently, the molecular mechanisms responsible for the remarkable cellular diversity of the *Atoh1* lineage are poorly understood. Furthermore, the origin of *Atoh1*- expressing cells, particularly those forming cerebellar granule cells – the largest population of neurons in the mammalian brain – is still an enigma. We have recently proposed that the posterior end of the cerebellar VZ, named as “posterior transitory zone (PTZ)”, contains bipotent progenitor cells for the *Atoh1* lineage and choroid plexus epithelium ^10^. However, the regulation and function of the PTZ have not been examined. A recent report showed that the RL in the developing human cerebellum has distinctive cytoarchitectural features from other vertebrates, including rhesus macaque ^11^. Unusual longevity of the human RL is attributed to the expansion of the posterior cerebellar vermis, a region that is associated with human cognition and predominantly affected in cerebellar birth defects such as Dandy-Walker malformation and cerebellar vermis hypoplasia ^11^. Therefore, studying the generation and maintenance of the PTZ will shed light on the evolution and congenital defects of the human cerebellum.

Conrad Waddington introduced the concept of epigenomic landscape to explain the emergence of distinct cell fates during development ^12^. In particular, the chromatin state defines the functional architecture of the genome by modulating the accessibility of *cis*-regulatory elements (CREs), which are regions of DNA bound by TFs to regulate the transcription of target genes. Together, the CREs and TFs constitute the regulatory logic for cell-state transition and lineage commitment. The adaptation of assay for transposase accessible chromatin (ATAC) allows measuring chromatin accessibility, a proxy for CRE activity, at single-cell resolution ^13^. Powerful algorithms have been developed to harness time-series single-cell data to identify the molecular trajectories that describe cell fate specification in vertebrate embryogenesis ^14–16^. Although several studies using single-cell technology have examined the heterogeneous cell populations in the mouse ^10, 17–21^ and human cerebellum ^22^, systematic integrated RNA and ATAC analysis by sampling developing cerebellar cells is still lacking.

In the current study, we applied single-cell RNA-sequencing (scRNA-seq) to resolve the developmental trajectories and the underlying transcriptional changes in the embryonic mouse cerebellum. We profiled the accessible chromatin of the mouse cerebellum between embryonic day (E) 12.5 and E14.5 using single-nucleus ATAC-seq (snATAC-seq). By integrating single-cell resolved RNA and ATAC modalities, we identified CREs that undergo temporal, cell-type-specific changes in chromatin accessibility and linked them to transcriptional targets. Based on the new information on the *trans-*, *cis-*elements and their linked targets, we reverse-engineered gene regulatory networks (GRNs) of individual cerebellar cell types and applied the GRNs to simulate genetic perturbations. Finally, we performed genetic and fate-mapping studies to validate our *in silico* analyses. Our study reveals the molecular control of the PTZ and its multipotency for granule neurons, Bergmann glia, and choroid plexus epithelium. Importantly, we show that the balance of cell fate specification of PTZ cell is crucial for the expansion of the posterior cerebellar vermis, a region the commonly affected in cerebellar birth defects in humans.

## RESULTS

### Reconstruction of cerebellar development with time-series scRNA-seq

To investigate developmental trajectories of cerebellar neural progenitor cells (NPC), we reanalyzed a previously published scRNA-seq dataset ^17^, which includes 12-time points from embryonic day (E) 10.5 to postnatal day (P) 10 mouse cerebella, focusing on the embryonic data (E10.5 - E17.5; Fig. S1A-B). Based on the known markers, we assigned identities to 29 recovered clusters, which included all major cerebellar cell types (Fig. 1A and Supplementary Table S1). In Uniform Manifold Approximation Projection (UMAP) embedding, proliferative cells, including NPC and granule cell progenitors (GCP), at either the synthesis (S) or the mitosis (M/G2) phase are arranged in circles, resembling the cell cycle, and linked to branches of postmitotic cells (Fig. 1B). Cells from different stages are well integrated but their relative contributions to each cell type change in agreement with the temporal development of the cell type (Fig. 1C-E) ^3^. For example, Purkinje cells (PC) initially appear at E12.5, GABAergic interneurons (IN) and GCP first appear at E14.5 and greatly expand afterward (Fig. 1D and E).

**Figure 1.**
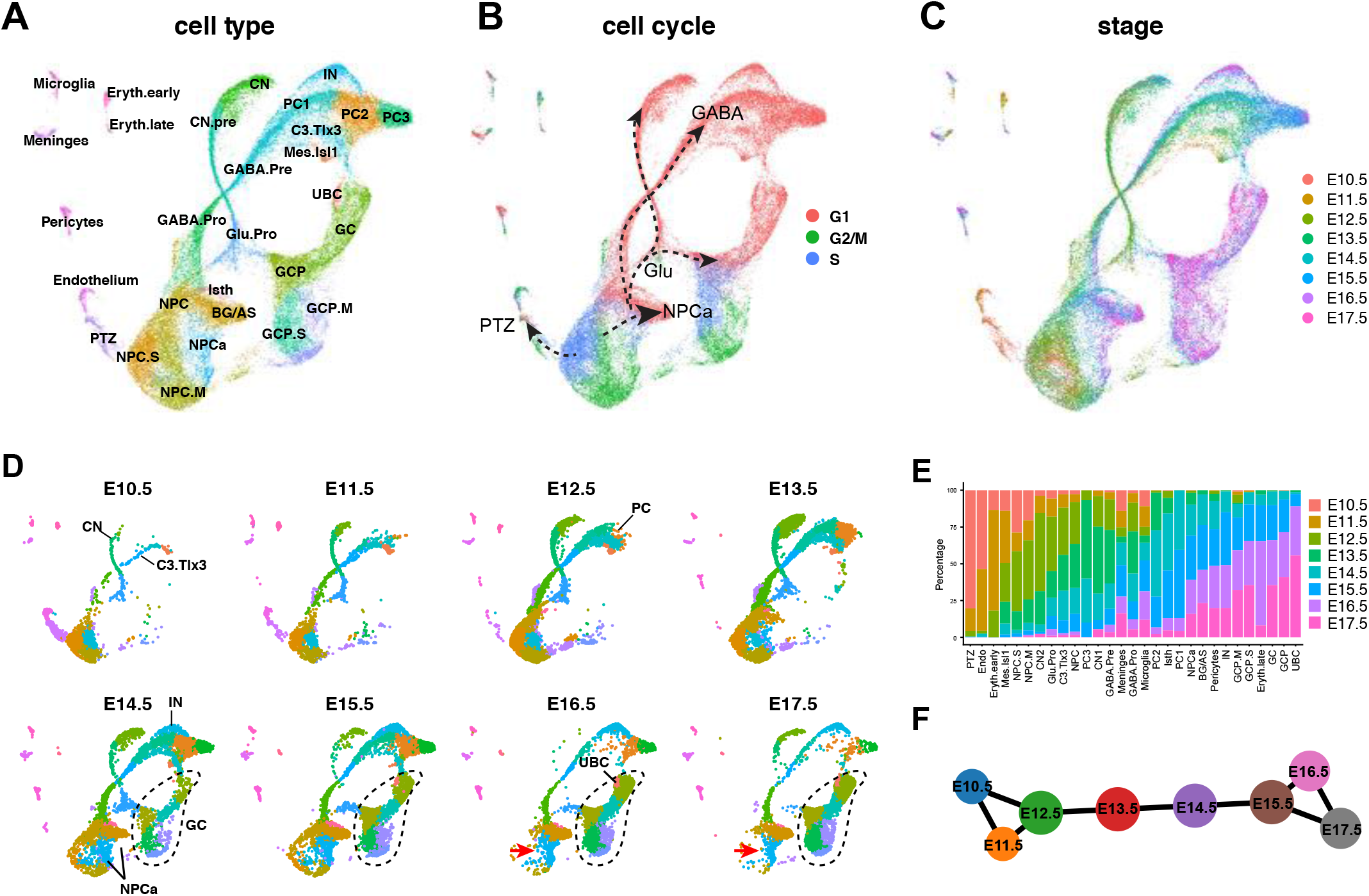
Reconstruction of the development of the embryonic mouse cerebellum by time-series scRNA-seq. (**A-D**) UMAP representation of merged scRNA-seq of mouse cerebella from E10.5 to E17.5. Dots indicate individual cells; colors show cell clusters (A,D), cell cycle phases (B), and embryonic stages (C); the dashed-line arrows in C denote the main branches of developmental trajectories arising from NPCs; the red arrow in D indicates an early-to-late shift of NPCs. The initial appearance of cerebellar nuclear neurons (CN), parabrachial nuclear neurons (C3.Tlx3), Purkinje cells (PC), interneurons (IN), cerebellar granule cells (GC), and unipolar brush cells (UBC) are indicated in D. Dashed lines in D circle GCs. (**E**) Bar plots showing cell composition across stages. (**F**) Partition-based graphic abstraction (PAGA) showing the relationship of cells from different stages.

The UMAP projection displays four main branches, corresponding to glutamatergic neurons, GABAergic neurons, posterior transitory zone (PTZ), and anterior neural progenitor (NPCa), respectively (Fig. 1B). After E13.5, NPCs shift toward the NPCa, displaying a transition from neural epithelium to radial glia with increasing expression of *Fabp7*, *Lxn*, *Aldh1l1*, and *Slc1a3* (Fig. 1C, D, and Supplementary Table S1). Along the glutamatergic branch, cells from the early stages form a continuum leading to cerebellar and extra-cerebellar nuclei (CN), whereas cells after E13.5 contribute to granule cells (GC; Fig. 1C-E), in agreement with a CN-to-GC switch in the production of glutamatergic cerebellar neurons ^5^. Therefore, scRNA-seq trajectory analyses reconstruct the order of cell differentiation in the embryonic mouse cerebellum.

### Resolution of four developmental trajectories from the cerebellar VZ progenitors

Unbiased hierarchical analysis using partition-based graph abstraction (PAGA) ^23^, correctly orders cells from E10.5 to E17.5 (Fig. S2A). Furthermore, PAGA reveals that cerebellar cells of the early (E10.5-E12.5) and late (E15.5-E17.5) are interconnected, whereas cells between E12.5 and E15.5 form a linear connection (Fig. 1F). This tripartite pattern suggests that the cell-state transition is dynamic and abrupt between E12.5 and E15.5, but it is much more gradual before or after that. We reasoned that, by taking advantage of the gradual and asynchronous development at the early phase, we could resolve the developmental trajectories from cerebellar NPC in greater detail. Indeed, UMAP embedding of E10.5-13.5 data reveals four conspicuous trajectories, each of which emerges in a step-wise manner (Fig. 2A and B). Inspecting selected genes revealed that these trajectories represent neurons arising from four cerebellar VZ domains that are demarcated by proneural genes *Atoh1*, *Ascl1*, *Ptf1a*, and *Neurog1*, respectively (Fig. 2C and D) ^24, 25^. The glutamatergic and GABAergic lineages are differentially marked by *Atoh1* and *Ascl1*; the latter gives rise to three branches, in which C2 and C4 are both marked by *Neurog1* while C2 is selectively marked by *Ptf1a* (Fig. 2C and D). C1 and C2 cells give rise to the cerebellar glutamatergic and GABAergic neurons, respectively ^24, 25^. C3 cells are precursors of the parabrachial nuclei ^26, 27^, whereas the fate of C4 cells is currently unknown. By performing immunofluorescence for the markers that represent each branch, we confirmed the cell types arising from the four cerebellar VZ domains (Fig. 2E-G).

**Figure 2.**
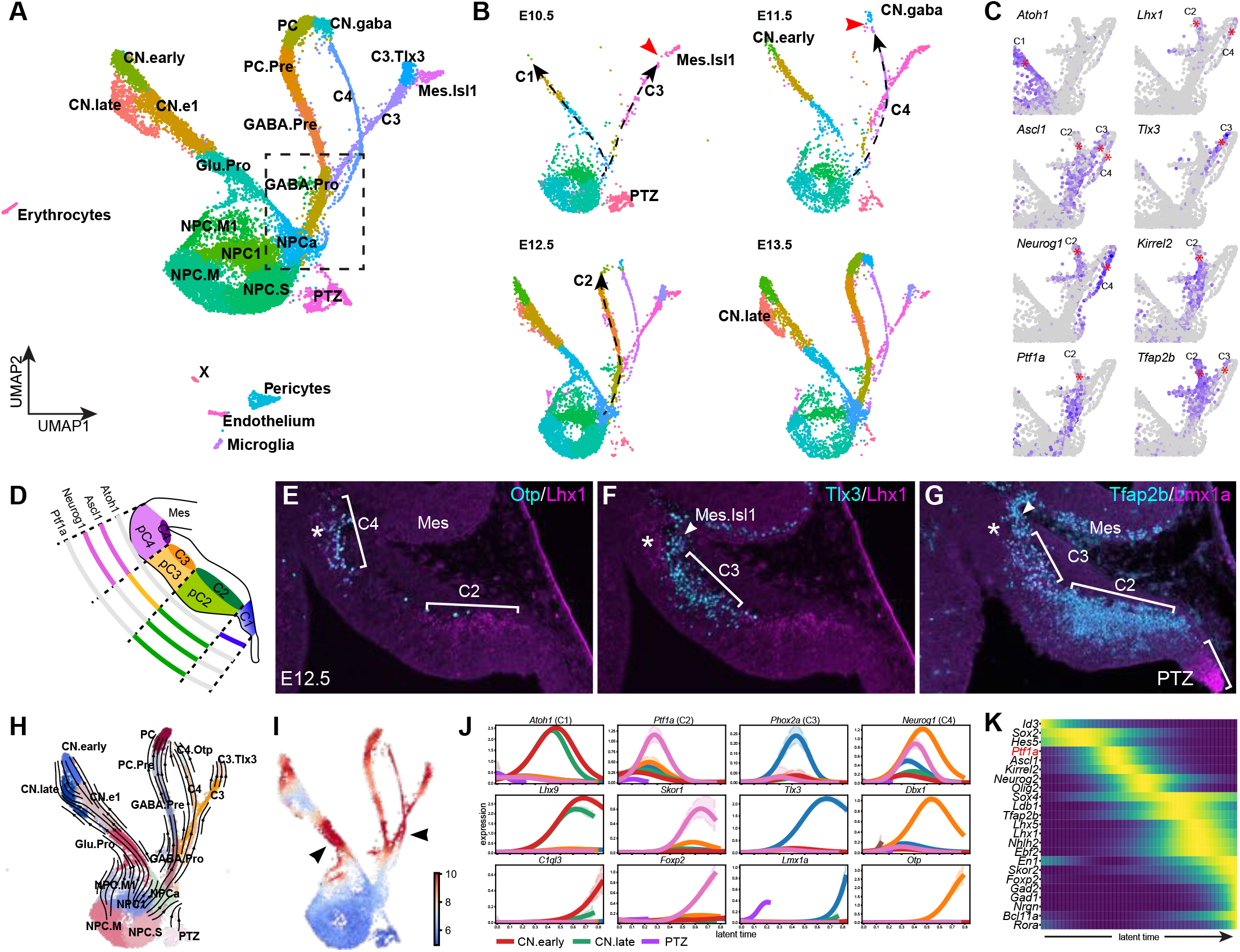
Cell fate choices during early cerebellar development. (**A** and **B**) UMAP projection of cells from the merged E10.5 - 13.5 dataset (A) and the individual stage (B). Dashed line arrows indicate the step-wise appearance of the trajectories; the arrowhead denotes the gap along the trajectory. (**C**) Expression of selected markers in the boxed UMAP area shown in A. The asterisk denotes the branch with the indicated gene. (**D**) Illustration of the regionalization of cerebellar VZ defined by the differential expression of proneural genes. (**E-G**) Immunofluorescence on sagittal sections of E12.5 mouse cerebella. Brackets demarcate C2-3 and the posterior transitional zone (PTZ); arrowheads indicate Isl1^+^ cells originated from the mesencephalon (Mes); asterisks denote C4 region, which is positive for Lhx1/5 but negative for Lmx1a, concordance with the expression pattern shown in C. (**H** and **I**) scVelo-inferred RNA velocity streamlines (H) and developmental kinetics (I) visualized in a UMAP embedding. Arrowheads indicate regions with accelerated differentiation. (**J**) Expression trends of selected genes across latent time for each developmental trajectory arising from cerebellar NPCs. (**K**) Heatmap showing the expression dynamics of regulators leading to the generation of PCs.

To study cellular differentiation kinetics, we calculated RNA velocity with scVelo ^28^. Adding RNA velocity vector streams to the UMAP embedding demonstrates the developmental trajectories – the proliferating state of NPCs traverses through the four branches (Fig. 2H). scVelo predicts that the differentiation progression is slow immediately after cell cycle exit and speeds up after cell fate commitment to the individual trajectory (Fig. 2I). Next, we identified cell fates, gene cascades, and driver genes along each developmental trajectory with CellRank (Fig. 2J, K, S2B, C, and Supplementary Table S2) ^29^. To validate the analysis, we performed functional enrichment analyses reasoning that genes in a given cascade should have common biological functions. Indeed, driver genes of each lineage exhibit distinct enrichments in biological function and pathways (Fig. S2D and E).

To assess the robustness of our results, we examined additional datasets. First, we repeated the analysis by adding E14.5 cells to the E10.5-13.5 dataset. We also reanalyzed other published scRNA-seq data of E10.5, E12.5, and E14.5 mouse cerebella ^21^. Finally, we generated our scRNA-seq data of E12.5, E13.5, and E14.5 cerebella (in-house). Remarkably, all these datasets produced almost identical results to the E10.5-13.5 one (Fig. S3). Collectively, our scRNA-seq analyses confirm the four subdomains in the cerebellar VZ in mouse embryos between E10.5 and E13.5. Importantly, we identify the genetic cascades, including putative driver genes, in the four populations of cells arising from the respective cerebellar VZ subdomains.

### Molecular control of cerebellar GABAergic neuron differentiation

Prior studies have demonstrated an essential role of *Ptf1a* in the production of GABAergic cerebellar neurons ^7^. Indeed, our computational analysis placed *Ptf1a* at the beginning of the gene cascade leading to the generation of PCs (Fig. 2K). We found that the expression of *Lhx1/5*, *Tfap2b*, and *Olig2,* which constitute the gene cascade downstream of *Ptf1a*, was absent from C2 in the E12.5 mouse cerebellum without *Ptf1a* (Fig. 3A-B’ and Fig. S4). Notably, the *Tlx3* expression, which normally demarcates the C3 area, was expanded into the C2 domain (Fig. 3A-B’ and S4), demonstrating that C2 cells are respecified to a C3 fate in the absence of *Ptf1a* as described previously ^27^. Although *Lhx1/5* and *Tfap2b* are expressed in both C2 and C4, their expression in C4 cells was unaffected in the *Ptf1a*-deficient cerebellum, indicating that C2 and C4 cells are differentially regulated. Therefore, our *in vivo* studies validate our single-cell trajectory inference and demonstrate the critical role of *Ptf1a* for the C2 cell fate.

**Figure 3.**
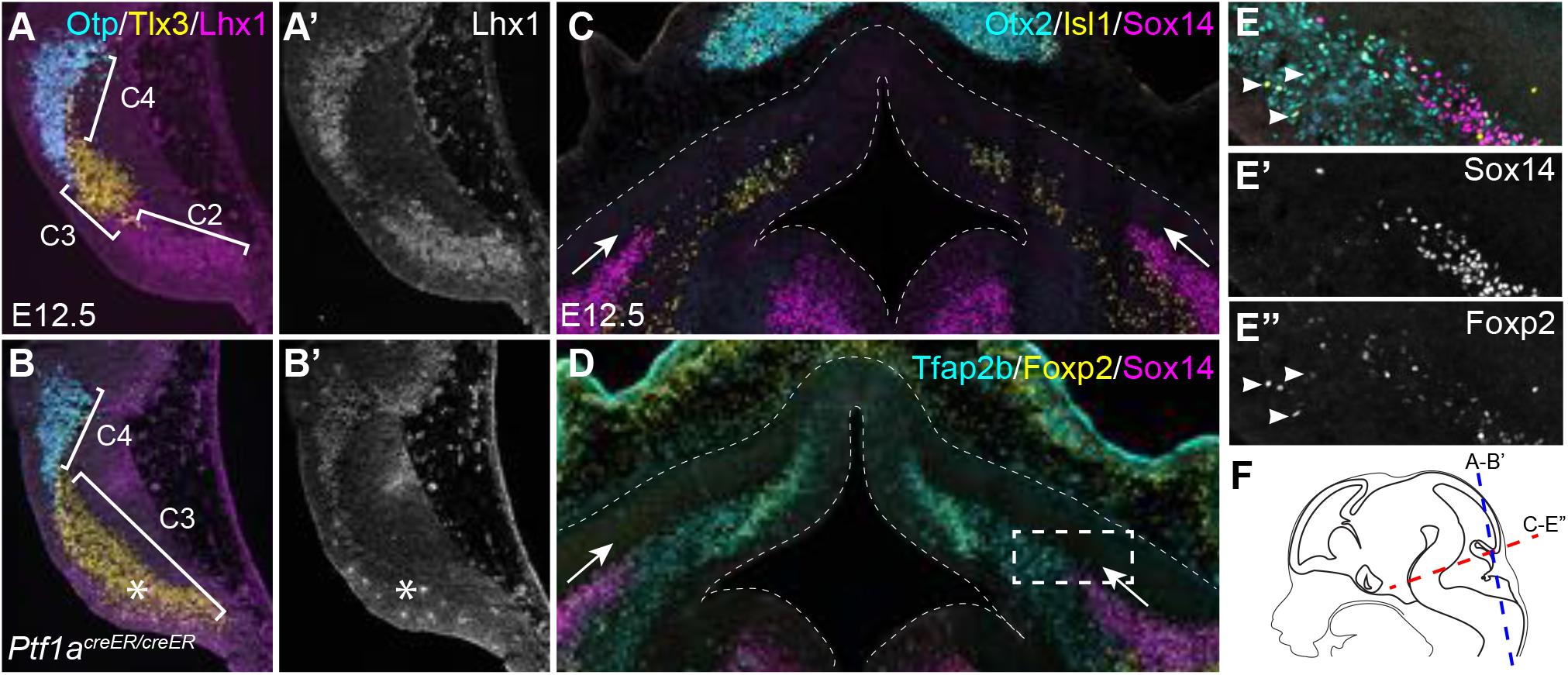
*In vivo* development of cerebellar GABAergic neurons. (**A-B’**) Immunofluorescence on coronal sections of wildtype and *Ptf1a*-deficient cerebellar anlage at E12.5. The asterisk indicates that the loss of Lhx1/5 accompanies an ectopic expression of Tlx3. Note that the anti-Lhx1 antibodies detect both Lhx1 and Lhx5. (**C-E”**) Immunofluorescence on horizontal sections of E12.5 cerebellar anlage. The boxed area in D is shown in separated channels in E-E”; dashed lines outline the cerebellar anlage; arrows indicate a ventral-to-dorsal extension of Sox14^+^ cells; arrowheads denote Foxp2^+^/Tfap2b^+^/Sox14^-^ cells, presumably nascent PCs, arising from the VZ. (**F**) Illustration showing the section planes.

In UMAP embeddings, cells are arranged according to the transcriptome similarity, but not necessarily the lineage relationship. We postulated that lineage-unrelated clusters could be detected by scrutinizing their temporal appearance along a trajectory. Indeed, although the Mes.Isl1 cluster, which represents mesencephalon-derived *Isl1*-expressing cells ^10^ resides at the end of the C3 trajectory, it is abundant and separated from the C3 branch at E10.5 (Fig. 2A and B), indicating that Mes.Isl1 and C3 cells are unrelated in the lineage. Prekop et al reported that *Sox14* marks GABAergic projection neurons in cerebellar nuclei ^30^. However, the origin of these neurons, denoted as CN.gaba, remains undetermined. Notably, CN.gaba cells first appear at E11.5 but they are separated from the C4 and C2 trajectories at this stage (Fig. 2B), suggesting that CN.gaba cells have a different origin from that of C2 or C4 cells. Indeed, Sox14^+^ cells were initially found in the lateral part of the cerebellar anlage, extending from the basal plate of r1 at E12.5 (Fig. 3C). Near the cerebellar VZ, newly-born neurons, presumably nascent PCs, were positive for both Foxp2 and Tfap2b, but negative for Sox14 (Fig. 3D-E”). Therefore, our data suggest that Sox14^+^ inhibitory projection neurons of cerebellar nuclei are originated from the basal plate of r1, rather than the cerebellar VZ.

### Ephemeral Atoh1 expressing cells destined for cerebellar nuclear neurons from the cerebellar VZ

The *Atoh1*-lineage arises from the RL and produces diverse cell types, including extra-cerebellar and cerebellar nuclear neurons, granule cells, and unipolar brush cells ^4, 5, 31^. Our data show that the early *Atoh1*-expressing cells produce early-born CN neurons (CN.early) before E12.5 and then late-born (CN.late) between E12.5 and E13.5 (Fig. 2A-C). CN.early likely represent isthmic nuclei neurons located in the anterior part of the nuclear transitory zone, whereas CN.late are cerebellar glutamatergic nuclear neurons ^10^. In contrast to the conventional view that GABAergic and glutamatergic neurons arise from spatially distinct germinal zones ^3^, our trajectory analysis suggests that the same group of NPCs gives rise to the *Atoh1*- and *Ascl1*- branches between E10.5 and E13.5 (Fig. 2A-C). To investigate the generation of the *Atoh1* lineage, we performed genetic fate mapping using an *Atoh1-Cre* transgene ^32^ and the Ai14 Cre-reporter mouse line, denoted as *R26^RFP^*, which expresses red fluorescent protein (RFP – tdTomato) upon Cre-mediated recombination ^33^. RFP-labeled descendants of the *Atoh1* lineage were found in the subpial stream ^34^, the mantle zone underneath the pial surface in *Atoh1- Cre;R26^RFP/+^* embryos at E10.5 and E11.5 (Fig. 4A-D’ and data not shown). Almost all RFP^+^ cells in the subpial stream were positive for Cre and Meis2 (99.4%; 335 RFP^+^ cells analyzed; Fig. 4B-D’). Surprisingly, radially oriented RFP^+^ cells were found in the dorsal and anterior part of the cerebellar VZ in addition to C1 (Fig. 4B-E). These RFP^+^ cells were positive for Sox2, but not Dcx, indicating that they were progenitors (Fig. 4F-G”). In contrast to the RFP^+^ cells in the subpial stream, only 35.1% of the RFP^+^ cells in the VZ were positive for Cre (262 RFP^+^ cells analyzed; Fig. 4B-D”). The expression pattern of RFP and Cre in the cerebellar VZ was undetected in the midbrain and telencephalon of *Atoh1-Cre;R26^RFP/+^* or in the cerebellum of *R26^RFP/+^* embryos (data not shown). These findings suggest that some NPCs in the cerebellar VZ transiently express Cre resulting in sporadic, but permanent, RFP labeling of their progeny. The fact that almost all RFP^+^ cells in the subpial stream were positive for Meis2 and Cre suggests that cerebellar VZ progenitors with ephemeral *Atoh1* expression are committed to a CN fate. In support of this notion, Cre^+^ or RFP^+^ cells were negative for Tfap2b and Ascl1, which mark precursors of GABAergic neurons (Fig. 4B-D’ and H). Collectively, our results suggest that some CN neurons are generated by NPCs that transiently express *Atoh1* in the cerebellar VZ.

**Figure 4.**
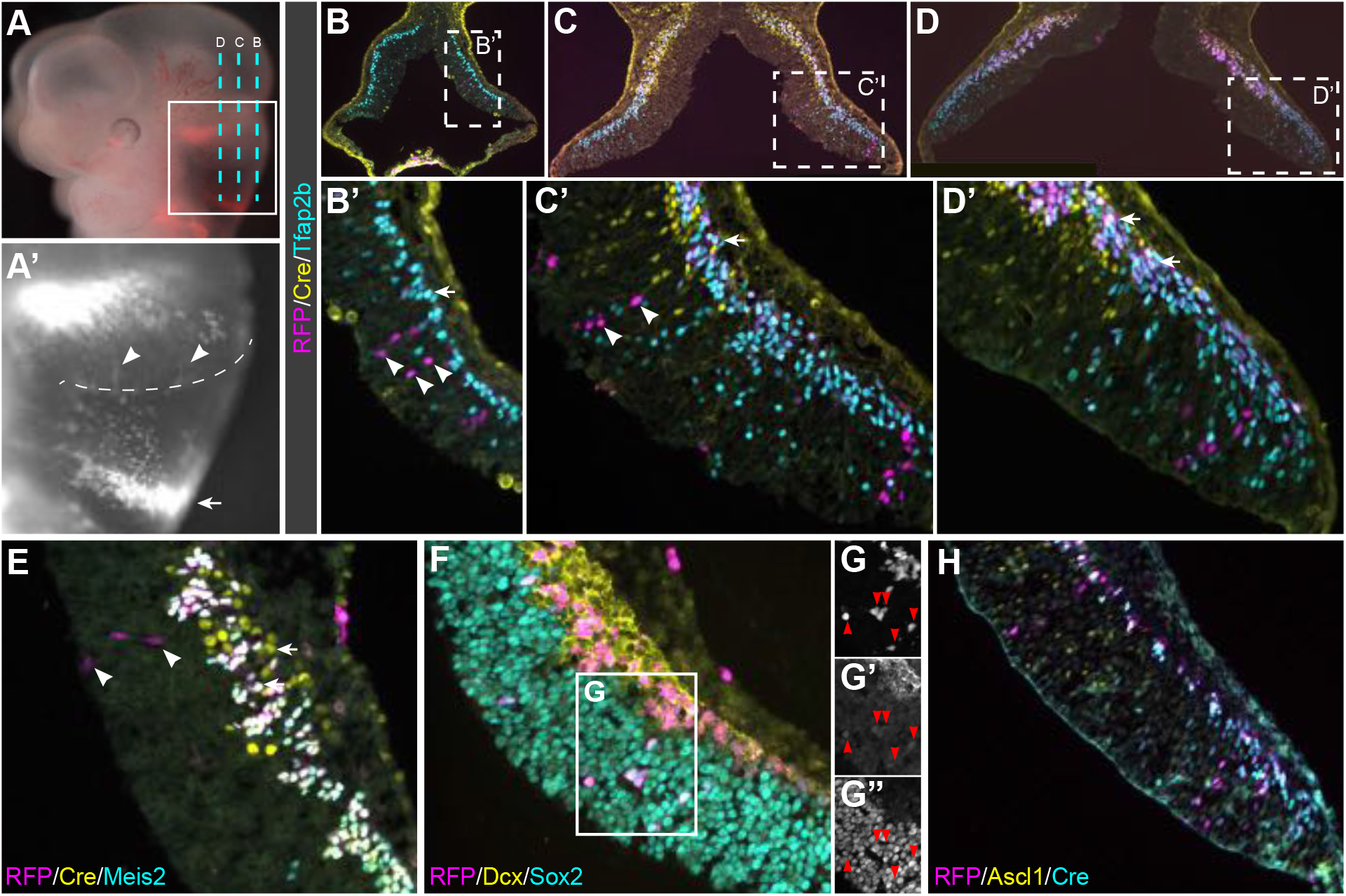
*Atoh1*-expressing cells arising from the cerebellar VZ. (**A, A’**) tdTomato fluorescence in wholemount *Atoh1-Cre; R26^RFP^*^/+^ mouse embryos at E10.5. The boxed area is enlarged in A’; the dashed lines indicate the section plane of B-D’ in A and the VZ-roof plate border in A’; arrowheads indicate RFP-labeled cells in the VZ. (**B-H**) Immunofluorescence of E10.5 *Atoh1-Cre; R26^RFP^*^/+^ cerebella. Arrowheads indicated radially oriented RFP^+^/Cre^-^ cells; arrows denote RFP^+^/Cre^+^ cells in the subpial stream.

### Single-cell analysis of chromatin accessibility of the embryonic mouse cerebellum

To identify CREs and quantify their dynamic activity during early cerebellar development, we performed snATAC-seq of the mouse cerebellum at E12.5, E13.5, and E14.5. Fragment size distribution and transcription start site enrichment demonstrated the high quality of our data (Fig. S5A-D). After stringent selection, we obtained a total of 31,144 high-quality cells, which were clustered into 26 groups (Fig. 5A-E and S5E). We determined fixed-width peaks (501bp) on aggregate single cells of the individual cell group and merged non-overlapping peaks to a union set of 401,835 peaks, which span over 200 Mb or 10.7% of the reference mouse genome.

**Figure 5.**
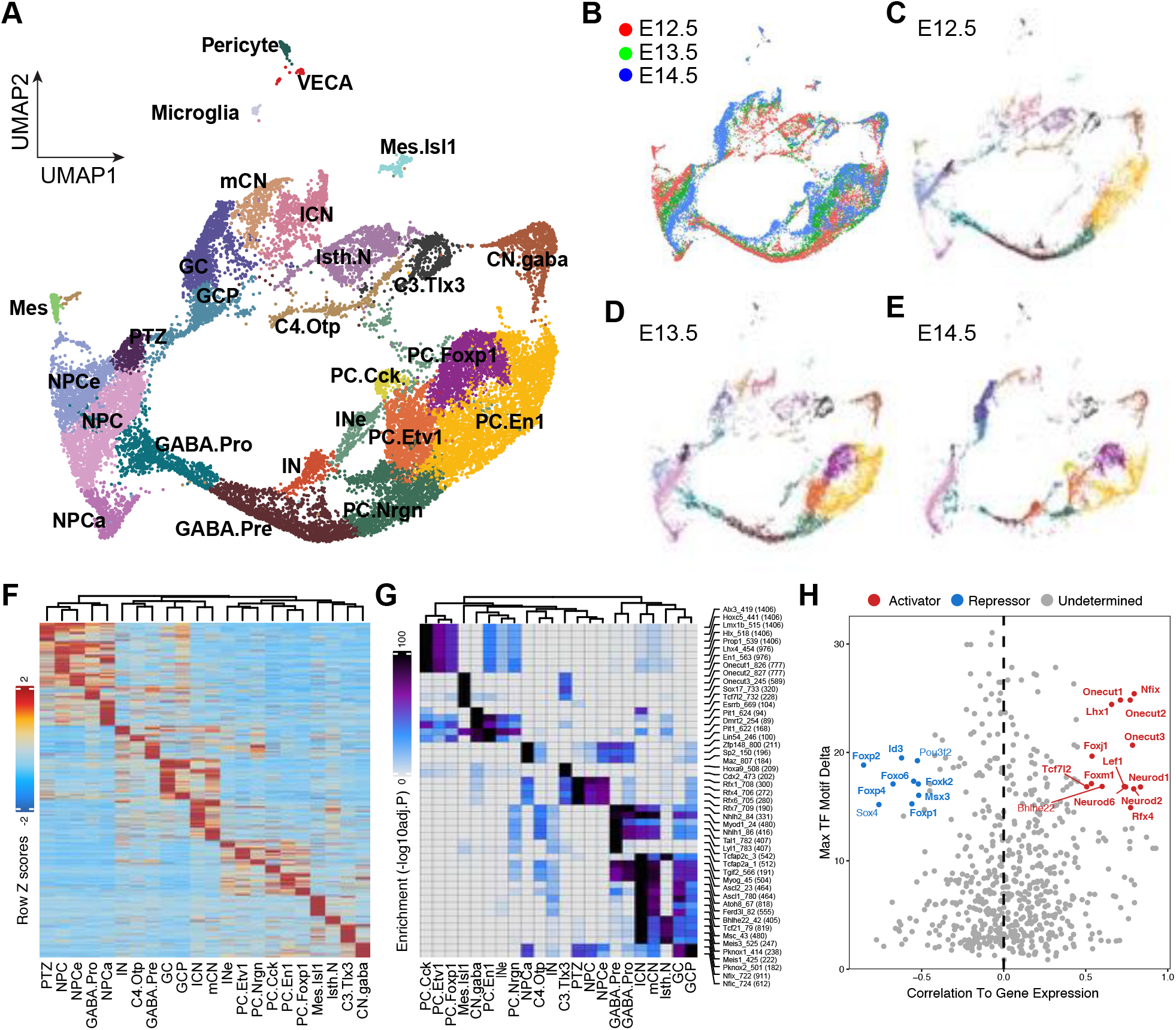
Studying the chromatin landscape in the embryonic mouse cerebellum through snATAC-seq. (**A-E**) UMAP projection of snATAC-seq cells. Colors indicate stages in B or cell types in the others. (**F** and **G**) Heatmap showing cluster-specific open chromatin regions (F) and the associated TF binding motifs (G). (**H**) Scatter plot showing the correlation between the expression of TFs and their binding motifs in the accessible chromatin region. Putative transcriptional activators and repressors are shown in red and blue, respectively.

To determine the identity of ATAC cell clusters, we calculated ‘gene activity scores’ to estimate gene expression bases on local accessibility of the gene body and promoter. We assigned identities according to the activity of known cell-specific markers: *Foxp2* and *Rora* for PC, *Ptf1a* for GABAergic progenitors (GABA.pro), *Sox14* for GABAergic CN projection neurons (CN.gaba), *Pax2* for GABAergic interneurons (IN), *Meis2* for glutamatergic CN neurons, *Pax6* and *Atoh1* for GC and GCP, respectively, and *Id3* for NPCs (Fig. S6A). Differential analyses identified cluster-specific markers and confirmed our identity assignment (Fig. S6B and Supplementary Table S3). To systematically assess the cell identity, we integrated our in-house E12.5-14.5 RNA and ATAC cells, revealing strong concordance between the two modalities – most ATAC cell clusters had one-to-one mapping to the RNA cell clusters (Fig. S6C). The integration resulted in a more accurate estimation of gene expression in each ATAC cell (Fig. S6D and E). Focusing on the E12.5 data, we showed that the global structures revealed by UMAP embedding for RNA and ATAC cells were remarkably conserved (Fig. S7A). When the scVelo-inferred latent time of the RNA cell was transferred to its nearest ATAC neighbors, we observed a smooth continuum of latent time in the chromatin manifold (Fig. S7B). One notable exception was that only the ATAC analysis correctly placed the PTZ cells between the NPC and the *Atoh1* lineage, even though both assays recovered the same set of molecular features of PTZ cells (Fig. S7C). The better performance of ATAC than RNA analysis in trajectory inference is likely because the former is unobscured by cell-cycle genes, which confound the RNA study of dividing cells ^35^.

### Examination of cell-type-specific accessible chromatin regions and their associated TF-binding motifs

We found that 40.5% of the total ATAC peaks exhibit differential accessibility among cell clusters (a median of 9,761 peaks per cluster; FDR < 0.01 and log2FC≥1; Fig. 5F), demonstrating that different cell types and states have distinctive chromatin landscapes. Motif analysis revealed that the cluster-specific loci are enriched for different TF binding motifs: the *Atoh1* lineage, including CN, GC, and GCP, is enriched for binding motifs of nuclear factor I TFs and basic helix-loop-helix TFs; NPCs for regulatory factor X TFs, and PCs for homeodomain-containing TFs (Fig. 5G). We curated a list of genomic features based on published snATAC-seq, bulk chromatin immunoprecipitation and DNAse I hypersensitive site (DNaseHS) sequencing to perform overlapping enrichment testing. As expected, cell-type specific open chromatin regions are largely consistent with those identified by a recent snATAC-seq of mouse cerebella (Fig. S6F) ^20^. Furthermore, the ATAC peaks specific for GABAergic progenitors are enriched for DNA sequenced bound by Ascl1 and Ptf1a ^36^, whereas those specific for GCs and GCPs are enriched for sequenced bound by Atoh1 (Fig. S6F) ^37^. Notably, chromatin regions identified by bulk-analysis of H3K27ac and DNaseHS of postnatal mouse cerebella are significantly enriched in GC-specific peaks, likely due to the prevalence of GCs in the postnatal cerebellum (Fig. S6F). Collectively, our finding demonstrate that different cerebellar cell types are regulated by distinct regulatory programs.

Next, we applied chromVAR ^38^ to calculate the relative motif activity at single-cell resolution. By correlating chromVAR deviation z-scores of TF motifs with expression levels of TFs, we systematically deduced putative positive and negative TFs in the developing cerebellum based on whether their expression was positively or negatively correlated with motif enrichment^39^. We iterated the analysis in stage-specific datasets and identified a total of 33 putative activators and 16 repressors (Fig. 5H). In support of this categorization, the majority of the TFs in these two groups are known transcriptional activators (29/33; 87.9%) or repressors (10/16; 62.5%), according to Gene Ontology Consortium (Supplementary Table S4). As the expression of these factors is closely correlated with the accessible chromatin region with their binding motifs, they may function as pioneer factors ^40^ and play crucial roles in cell fate specification in the developing cerebellum.

### Identification of cis-regulatory elements for cell-type-specific transcriptional programs

To find CREs, we identified chromatin accessible peaks that correlated with transcription in the ATAC-RNA joined cell clusters. We iterated the analysis for individual stages and identified a total of 44,329 peak-to-gene pairs with 31,088 unique accessible peaks and 7,159 target genes. 69.4% of the identified CRE-target pairs are specific to a particular stage, demonstrating the temporally dynamic activation of most CREs. The median distance between CREs and target promoters is 17,1101 kb, 14.9 times greater than that to the closest genes (Fig. 6A). Most CREs are located in the intronic or distal intergenic regions (Fig. 6B), suggesting that the identified CREs are mostly distal elements. While 73.0% of the CREs are assigned to a single gene, the rest are linked to two or more (up to 13; Fig. 6C), showing that some CREs may regulate multiple genes. The median number of CREs per gene is four (Fig. 6D). Remarkably, the top 10 percentile genes with the highest numbers of CREs (>16) are enriched for DNA-binding transcription regulator activity and regulation of neural development (Fig. 6E, F, and Supplementary Table S5). Among the top eight genes, *Zic5, Zic2, Pax6, Hes5, Sox9, Lhx1, Sox2,* and *Pax3*, are well-known TFs important for cerebellar development. These observations suggest that key TFs acting in early cerebellar development are subjected to complex regulation via numerous CREs in agreement with a recent report ^20^.

**Figure 6.**
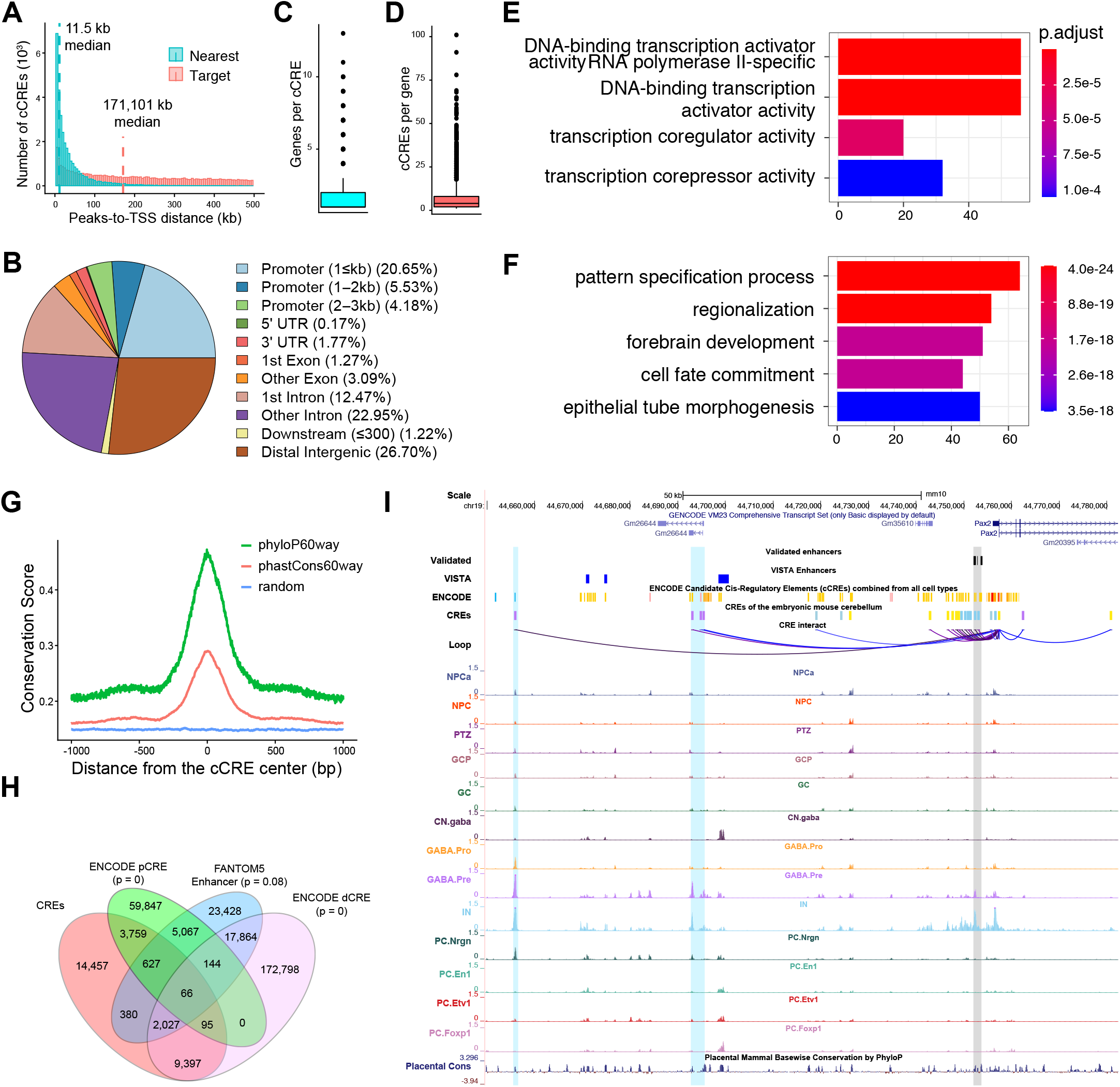
Characterization of *cis*-regulatory elements in the embryonic mouse cerebellum. (**A**) Histograms comparing the distribution of distances from CREs to the nearest genes versus target genes. (**B**) Pie chart showing the distribution of CREs among genomic features. (**C** and **D**) Box-Whisker plots showing the number of genes per CRE (C) and the number of CREs per gene (D). (**E** and **F**) Bar plots showing enrichment of Gene Ontology in molecular function (E) and biological process (F). (**G**) Distribution of the average based-pair evolutionary conservation scores using 60-vertebrate genome alignments at regions around CREs and randomly shuffled regions. (**H**) Venn diagram showing overlapping of CREs with proximal- (pCRE) and distal-elements (dCRE) identified by ENCODE or enhancers by FANTOM5. The percentages indicate the elements unique to each group. (**I**) Examples of linkages of CREs and targets. The grey areas highlight the CRE that has been validated by transgenic mouse assays.

Additional lines of evidence support the functional significance of the identified CREs. First, we detected a sharp increase in evolutionary conservation scores at the center of CREs (Fig. 6G). Furthermore, the CREs are significantly overlapped with putative enhancers identified by the ENCODE consortium and FANTOM5 (Fig. 6H) ^41, 42^. Finally, we asked whether our approach recovered the enhancers that had been validated by transgenic mouse assays. We compiled a list of the enhancers of seven genes that are involved in cerebellar development, including *Atoh1*, *Fgf8*, *Kirrel2*, *Pax2*, *Pcp2*, *Ptf1a*, and *Wnt1* ^43–49^. Remarkably, all these validated enhancers were recovered by our study (Fig. 6I and S8). Therefore, our integrative scRNA-seq and snATAC-seq analysis provide enriched information on the CREs that control the temporospatial gene expression in the developing cerebellum.

### Reconstruction of gene regulatory networks governing cerebellar development

We leveraged the newly identified CREs to reverse-engineer GRNs by applying CellOracle ^50^. For each cell type and state at E12.5 (Fig. 7A), we produced a GRN, which exhibits a scale-free network characteristic for biological networks (data not shown). The network entropy, indicative of the average undifferentiated state, decreases from NPC to PC or from GCP to CN as expected (Fig. 7B). We identified key regulators of each cell-specific GRN through network analyses (Fig. 7C, S9, and Supplementary Table S6). Among the predicted top regulators of GABAergic progenitors, *Neurog1/2*, *Ptf1a*, *Ascl1*, *Olig2/3*, *Tfap2a/b*, and *Lhx1/5* have been shown to play crucial roles in the development of GABAergic neurons in the cerebellum ^7, 51–56^. Similarly, *in vivo* studies have demonstrated the essential role of *Atoh1*, *Neurod2*, *Barhl1*, and *Lhx9* in cerebellar glutamatergic neurons as predicted by our GRN analysis (Fig. 7C) ^6, 57–60^. In the GRN of GABAergic progenitors, the majority of the CREs of the inferred targets of Ascl1 and Ptf1a are bound by Ascl1 (76.7%) and Ptf1a (75.0%) according to a prior binding profile study (Fig. 7D and E) ^36^.

**Figure 7.**
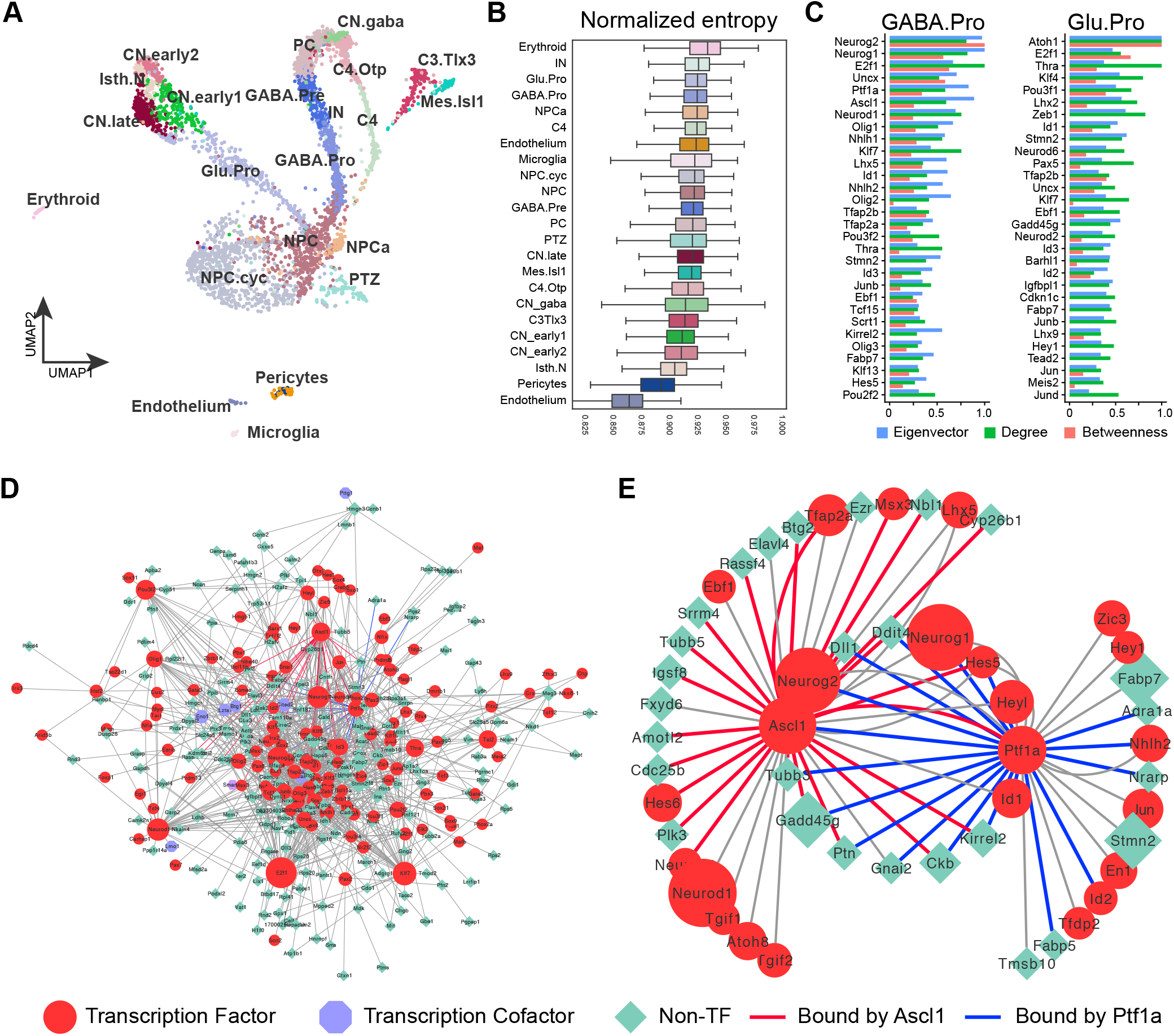
Reconstruction of gene regulatory networks of cerebellar cell types and states. (**A**) UMAP projection of in-house E12.5 scRNA-seq. (**B**) Box plot showing the distribution of network entropy scores for each cell cluster. (**C**) Top 30 driver genes by an aggregate ranking of degree centrality, betweenness centrality, and eigenvector centrality in the GRNs of GABAergic (GABA.Pro) and glutamatergic progenitor (Glu.Pro). (**D**) The inferred GRN of GABA.Pro cells. (**E**) Subnetworks of D with all targets of Ascl1 and Ptf1a. Links confirmed by ChIP-seq are shown in red (for Ascl1) and blue (for Ptf1a).

### Studying the molecular regulation of PTZ development via GRN simulations

To evaluate the validity and utility of the GRNs, we applied CellOracle to simulate how TF perturbations affect cell fate decisions. We focused on *Ptf1a* and *Atoh1*, two well-characterized master regulators in cerebellar development. In agreement with mouse genetic studies ^6, 7, 31^, GRN simulations predicted that the loss of *Ptf1a* would reduce GABAergic progenitors, whereas the loss of *Atoh1* would deplete glutamatergic progenitors (Fig. S10A and B). We also simulated *Ptf1a^Atoh^*^1^ and *Atoh1^Ptf1a^* mutations, two reciprocal knock-in alleles ^9^. In concordance with the *in vivo* outcome ^9^, simulations indicated that *Ptf1a^Atoh1^* increased glutamatergic progenitors at the expense of GABAergic progenitors, whereas *Atoh1^Ptf1a^* produced the opposite effect (Fig. S10C and D). Unexpectedly, GRN simulations suggested that the loss of *Ptf1a* would enlarge the PTZ (Fig. 8A and B). Based on the overlapping expression pattern of Atoh1, Lmx1a, Hes1, Otx2, Pax6, and Reln (Fig. S7C), we identified the PTZ in the wildtype cerebellum and confirmed its enlargement in *Ptf1a*-deficient cerebellum at E13.5 (Fig. 8C-E).

**Figure 8.**
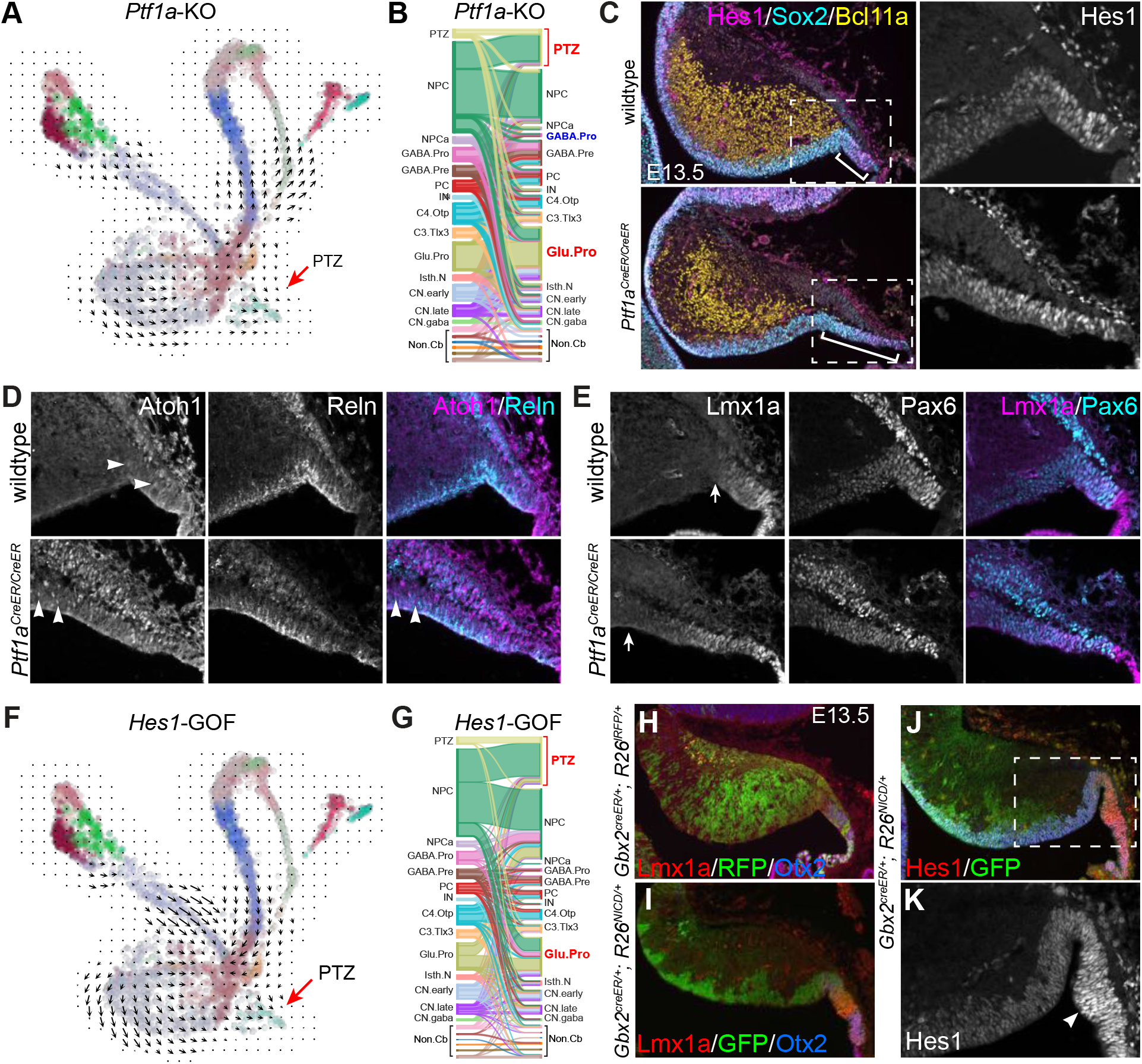
Investigation of the molecular control of PTZ development by GRN simulations and genetic experiments. (**A, B**) The vector field graph and Sankey diagram showing simulated cell transition caused by the loss of *Ptf1a*. The arrow and bracket indicate the enlarged PTZ. (**C-E**) Immunofluorescence on sagittal sections of E13.5 cerebella. The boxed areas are enlarged in D and E; the brackets indicate the enlarged PTZ domain, which is marked by *Hes1*, *Reln*, and *Lmx1a*; arrowheads indicate nascent Atoh1^+^ cells. (**F,G**) The vector field graph and Sankey diagram showing the effects of *Hes1* over-expression. (**H-K**) Immunofluorescence on cerebellar sagittal sections of E13.5 *Gbx2^CreER/+^;R26^RFP/+^* or *Gbx2^CreER/+^;R26^NICD/+^* mice that were given tamoxifen at E8.5. The boxed area in J is enlarged in K; the arrowhead shows robust Hes1 expression in the enlarged PTZ.

*Hes1* and *Hes5,* which are important for NPC maintenance ^61, 62^, are expressed in the cerebellar VZ, whereas only *Hes1* is expressed in the PTZ (Fig. S11A and B). GRN simulations predicted that the loss or gain of function of *Hes1* would reduce or enlarge the PTZ, respectively (Fig. 8F, G, and S11C). *Hes1* is a well-known transcriptional target of Notch signaling ^63^. Upon activation, the Notch intracellular domain (NICD) is released from the cell membrane and translocated into the nucleus to promote *Hes1* transcription ^63^. We predicted that an ectopic expression of *NICD* would induce *Hes1* resulting in a gain of function (GOF) of *Hes1*. To this end, we conditionally expressed *NICD* in the cerebellar VZ by combining *Gbx2^CreER^* ^64^ and *R26^NICD^*, in which *NICD-ires-GFP* is expressed from the *Rosa26* locus upon Cre-mediated recombination ^65^. After tamoxifen administration between E8.5 and E10.5, we detected descendants of the *Gbx2*-lineage throughout the cerebellum in *Gbx2^CreER/+^;R26^RFP/+^* embryos at E13.5 (Fig. 8H). By contrast, *NICD*-expressing (*NICD^+^*) cells, which were marked by GFP-immunoreactivity, were restricted to the VZ and PTZ of *Gbx2^CreRE/+^;R26^NICD/+^* embryos (Fig. 8I and J). As expected, robust *Hes1* expression was detected in *NICD^+^* cells, particularly in the PTZ (Fig. 8J and K). The PTZ was markedly enlarged in *NICD*-GOF embryos at E13.5 (Fig. 8H-K), demonstrating the positive regulatory role of *Hes1* in PTZ development as predicted by GRN stimulations. Therefore, we have revealed the crucial role of *Hes1* in PTZ development while demonstrating the efficacy of GRN stimulations to predict the outcome of TF perturbations.

### Important roles of the PTZ in the growth of the posterior cerebellar vermis

In contrast to the sparse contribution of progenies of *Gbx2*-expressing cells labeled at E9.5 to the choroid plexus in the control, abundant *NICD*^+^ cells were found in the exceedingly enlarged choroid plexus in E16.5 *Gbx2^CreRE/+^;R26^NICD/+^* embryos (Fig. 9A-C). Although RFP- labeled descendants of the *Gbx2* lineage abundantly contribute to the EGL, few *NICD*^+^ cells were detected in the EGL (Fig. 9B and C). Within the PTZ, the expression of NICD (GFP) and Atoh1 was mutually exclusive, indicating a cell-autonomous inhibition of *Atoh1* by NICD (Fig. 9D-D”). A gap was detected between the EGL and PTZ in E16.5 *Gbx2^creRE/+^;R26^NICD/+^* embryos that were given tamoxifen at E8.5 or E9.5, suggesting that PTZ cells fail to replenish the *Atoh1* lineage in the presence of Notch signaling (Fig. 9B and C). In the cerebellar cortex, almost all *NICD*^+^ cells displayed the morphology and molecular features characteristic for Bergmann glia, including the expression of Fabp7, Mki67, Sox2, and Sox9 ^66^, indicating that Notch promotes Bergmann glia generation (Fig. 9E and F).

**Figure 9.**
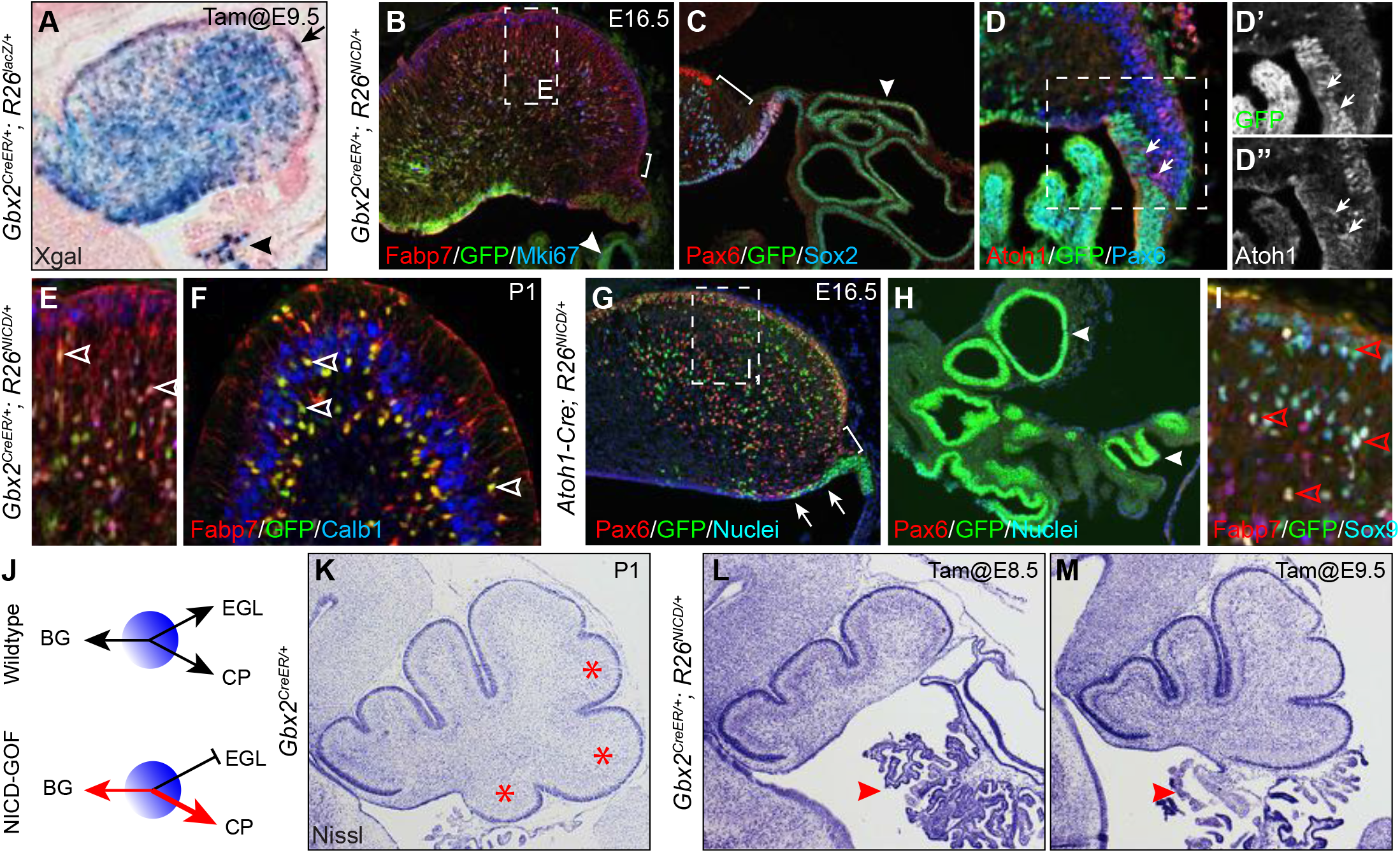
Specific loss of the posterior cerebellar vermis due to the abnormal cell-fate decision in the PTZ. (**A**) Xgal histochemistry showing the ubiquitous distribution of *Gbx2*- expressing cells labeled at E9.5 in the E16.5 cerebellum. The arrow indicates Xgal+ cells in the EGL; the arrowhead shows a few Xgal^+^ cells in the choroid plexus (CP). (**B-F**) Immunofluorescence on cerebellar sagittal sections of *Gbx2^CreER/+^;R26^NICD/+^* mice that were given tamoxifen at E10.5. The brackets indicate the discontinuous EGL; arrowheads show the accumulated NICD^+^ cells in the enlarged CP; arrows show Atoh1^+^ cells devoid of NICD; the empty arrowhead denotes NICD^+^ Bergmann glia. (**G-I**) Immunofluorescence on cerebellar sagittal sections of *Atoh1-Cre;R26^NICD/+^* mice at E16.5. Arrowheads show NICD^+^ cells accumulated in the expanded CP; arrows indicate *NICD*^+^ cells in the VZ outside the PTZ; the empty arrowhead denotes NICD^+^ Bergmann glia. (**J**) Illustration of how Notch regulates the tri-potency of PTZ progenitors. (**K-M**) Nissl histology of sagittal section of P1 cerebellum. The arrowheads indicate the CP that is greatly expanded. Note the progressive loss of posterior cerebellar vermis (asterisk in K) when NICD was induced *Gbx2^CreER/+^;R26^NICD/+^* at E8.5 and E9.5.

As *NICD* was induced broadly in the cerebellar VZ of *Gbx2^creRE/+^;R26^NICD/+^* cerebella, it was unclear whether Notch activation induced Bergmann glia from the PTZ. To address this question, we repeated the *NICD*-GOF experiment with *Atoh1-Cre*. As expected, RFP^+^ fate-mapped *Atoh1* progenies were predominantly found in the EGL of *Atoh1-Cre;R26^RFP/+^* cerebellum at E18.5 (Fig. S12A and A’). Although a few RFP^+^ cells were present in the choroid plexus, none became Bergmann glia (Fig. S12A and A”). When *NICD* was induced in the *Atoh1* lineage, many *NICD*^+^ cells were detected in granule cells and cerebellar nuclear neurons of *Atoh1-Cre;R26^NICD/+^* embryos (Fig. 9G, S12B, and C). However, *NICD^+^* cells were hardly detected among nascent Atoh1^+^ cells leaving the PTZ and a gap was detected between the PTZ and EGL in the *Atoh1-Cre;R26^NICD/+^* cerebellum as that found in *Gbx2^creRE/+^;R26^NICD/+^*, conforming the inhibitory role of Notch in *Atoh1* induction (Fig. 9C, G, and S12B). These findings show that GCPs tolerate persistent *NICD* expression and therefore, the lack of *NICD^+^* cells in the EGL of *Gbx2^creRE/+^;R26^NICD/+^* embryos results from abnormal cell-fate specification rather than selective cell loss. In contrast to the almost exclusive contribution of RFP^+^ cells to glutamatergic neurons in *Atoh1-Cre;R26^RFP/+^* embryos, abundant *NICD^+^* cells were found in the exceedingly expanded choroid plexus and Bergmann glia in *Atoh1-Cre;R26^NICD/+^* embryos (Fig. 9H and J). Therefore, the forced expression of *NICD* in nascent *Atoh1*-expressing cells respecifies the *Atoh1* lineage to form choroid plexus epithelium and Bergmann glia.

*Wnt1* expressing cells contribute to the EGL, choroid plexus, and occasionally Bergmann glia ^67^. In *Wnt1-CreER;R26^NICD/+^* embryos that received tamoxifen at E8.5, *NICD^+^* cells were accumulated in the greatly enlarged choroid plexus, but hardly found in the EGL (Fig. S12D and E). We detected abundant *NICD^+^* cells that co-expressed Fabp7 and Sox9 and displayed typical Bergmann glial morphology (Fig. S12D and E). Collectively, our data show that the activation of Notch signaling promotes PTZ progenitors to form choroid plexus epithelium and Bergmann glia at the expense of the *Atoh1* lineage. Therefore, the PTZ represents a dynamic germinal zone containing multipotent progenitors for Bergmann glia, glutamatergic neurons, and choroid plexus (Fig. 9J).

In *Gbx2^CreRE/+^;R26^NICD/+^* mice that were given tamoxifen between E8.5 and E9.5, the posterior part of the cerebellum was truncated while the choroid plexus was greatly expanded (Fig. 9K-M). A similar phenotype was found in *Wnt1-CreER;R26^NICD/+^* cerebellum at P15 (Fig. S12F-H). The cerebellar hypoplasia was more severe near the midline than the lateral region (Fig. S12G, H, and data not shown). Our results demonstrate that the multipotent progenitor cells in the PTZ replenish *de novo Atoh1* lineage to sustain the growth of the posterior part of the cerebellar vermis.

## DISCUSSION

### Integrated analyses of transcriptome and chromatin accessibility at single-cell resolution

Through scRNA-seq trajectory analyses, we show the developmental dynamics of the entire cerebellar primordium and in individual lineages during mouse embryogenesis (Fig. 1F and 2A-I). We confirm the four different cellular populations originating from distinct compartments of the cerebellar VZ and infer gene expression cascades along their developmental trajectories (Fig. 2 and S2). Our results provide new information on the molecular mechanisms that create the heterogeneity of glutamatergic (see later) and GABAergic neurons. We suggest that Sox14^+^ inhibitory projection neurons of the cerebellar nuclei may be originated outside the cerebellar primordium, probably from the basal plate of the neural tube. Together with our previous finding of mesencephalon-derived Isl1^+^ cells ^10^, we show that the migration of cells originated outside the cerebellar anlage may represent an important mechanism to increase the cellular diversity of the cerebellum.

By applying paired single-cell RNA-ATAC analysis, we have systematically studied the regulome that governs cell state transition and lineage commitment in the developing mouse cerebellum. We identify 31,088 candidate CREs with their targets and predicted transcription factors that act through these candidate CREs. The reference maps of CREs for the mouse cerebellum would not only help to understand the mechanism of gene regulation in different cell types but also enable targeting and purifying of specific cell types. To this end, we strive to make our data broadly available to the community (see Data Availability).

Despite the striking capability of GRN analysis, there are limitations and dependencies. First, the accuracy of the predicted TFs based on peak-gene pairs depends on the availability of high-quality binding motifs. Second, when simulating TF perturbation, CellOracle predicts the directionality of the fate changes in affected cells expressing the particular TF, but not the long-term consequences. Furthermore, only TFs with relatively high variability and expression values are available for perturbation simulations.

### The cerebellar VZ contains ephemeral Atoh1-expressing progenitors destinated to cerebellar nuclear neurons

Past studies have demonstrated the extraordinary diversity of *Atoh1* derivatives in the cerebellum ^4, 5, 31^. *Atoh1*-expressing cells arising from the PTZ between E9.5 and E12.5 produce cerebellar nuclear (CN) neurons ^5^. It has been shown that the earliest neural precursor cells exit the cell cycle from the cerebellar VZ at E10.25 and contribute to CN neurons ^68, 69^. These early-born CN neurons express *Irx3*, *Meis2*, and *Lhx2/9,* and they undergo radial migration to reach the nuclear transitory zone ^68^. The relationship between the early-born CN neurons from the VZ and the *Atoh1* lineage has never been determined. Here, we provide evidence that the early-born CN neurons present by E10.5 are probably derived from *Atoh1^+^* cells (Fig. 2A and B). Furthermore, NPCs with ephemeral *Atoh1* expression give rise to CN neurons at E10.5 (Fig. 4 and see next section). Therefore, our observations reconcile the previous findings on the origin of CN neurons. Green et al, showed that some *Atoh1*-expressing cells are generated at the isthmus, independent of the PTZ, producing isthmic nuclei ^70^. Future studies should determine whether the early-born CN neurons contribute to the isthmic nuclei.

NPCs, including those in the anterior part of the cerebellar VZ, express Cre that is driven by an *Atoh1* enhancer (Fig. 4). In contrast to the relative abundance of Cre^+^ cells, RFP^+^ fate-mapped cells arising from the VZ are rare and mostly negative for Cre in *Atoh1-cre;R26^RFP/+^* embryos, suggesting that a small percentage of Cre^+^ NPCs is labeled due to the ephemerality of Cre expression. Importantly, almost all RFP^+^ cells are negative for Tfap2b and Ascl1, and become positive for Meis2 and Cre when they reach the mantle zone (Fig. 4), indicating that the NPCs that transiently express *Atoh1* are committed to CN neurons. Basic helix-loop-helix transcriptions, like Ascl1, Hes1, and Olig2, are expressed in an oscillation manner to stabilize the stemness of NPCs ^71, 72^. Future studies should determine whether oscillatory control of *Atoh1* specifies and sustains CN progenitors in the cerebellar VZ.

### The posterior transitory zone contains multipotent progenitor cells

Although the RL is commonly referred to as the origin of cerebellar glutamatergic neurons, there is no consensus on the exact definition of the RL. Gene expression studies suggested that there are four molecularly distinct RL compartments ^73^. However, we were unable to resolve discrete cell groups corresponding to the presumed compartments through scRNA-seq^10^. Instead, we identified an inseparable NPC cluster that possesses the potential to form both the *Atoh1* lineage and choroid plexus epithelium and mapped the cell group to a triangular structure interfacing the VZ, choroid plexus, and the emerging EGL ^10^. We named this structure the posterior transitory zone (PTZ) as we postulated that cerebellar NPCs are actively recruited into the PTZ to replenish progenitors destined for glutamatergic neurons and choroid plexus epithelium ^10^. In the present study, we refine the molecular feature of PTZ cells (Fig. S2D,E, and S7C). We show that *Hes1* is expressed in the PTZ, which specifically lacks *Hes5* (Fig. S11B). Through *in silico* and *in vivo* studies, we demonstrate that the elevated expression of *Hes1*, as a result of the loss of *Ptf1a* or forced expression of *NICD*, enlarges the PTZ, suggesting that *Hes1* is important to recruit NPCs and maintain PTZ cells (Fig. 8).

Notch activation promotes PTZ cells to form choroid plexus epithelium but inhibits *Atoh1* expression (Fig. 9). The latter is in agreement with the previous report that Notch activation inhibits *Atoh1* expression by antagonizing BMP signaling ^74^. Somewhat unexpectedly, we found that the forced expression of *NICD* induced Bergmann glia not only from *Gbx2*- expressing cells but also from the *Atoh1* lineage, demonstrating that Notch activation respecifies nascent *Atoh1* cells to form Bergmann glia and choroid plexus from glutamatergic neurons. Although it is well known that Notch signaling is important for Bergmann glia differentiation ^75^, our results demonstrate the involvement of Notch signaling in the induction of Bergmann glia. We have previously shown that an FGF-ERK-ETV axis is essential for the transition of cerebellar radial glia to Bergmann glia ^66, 76^. How Notch interacts with the FGF-ERK-ETV axis to induce Bergmann glia awaits to be examined. Prior studies have shown that nascent Bergmann glia arise from a VZ domain, the so-called “peritrigonal glial matrix” ^77^, immediately anterior to the PTZ ^76, 78, 79^. The progenies of *Wnt1-*expressing progenitors, which are confined to the PTZ (Fig. S7C), contribute to glutamatergic neurons, choroid plexus, and Bergmann glia ^67^. These observations suggest that precursors of Bergmann glia must have a labile cell fate interchangeable with that of the PTZ. Altogether, our results demonstrate that the PTZ represents a highly dynamic stem cell zone, where the balance of cell fate commitments is regulated by Notch signaling, in part through Hes1 (Fig. 9J).

### The maintenance and differentiation of PTZ cells control the expansion of the posterior cerebellar vermis

Disproportionate reduction in the posterior cerebellum is a hallmark of many human cerebellar congenital defects, including Dandy-Walker malformation and cerebellar vermis hypoplasia ^11^. It was suggested that spatiotemporal expansion of the RL may be specific to humans, raising the question of whether rodents are suitable to study cerebellar malformations in humans ^11^. In the present study, we demonstrate that the active recruitment of NPCs into the PTZ to replenish the *Atoh1* lineage is essential for the enlargement of the posterior cerebellar vermis. Interestingly, the *NICD*-GOF mice display features that clinically define Dandy-Walker malformation, including a hypoplastic, upwardly rotations vermis, an enlarged fourth ventricle, and an enlarged posterior fossa ^80^. Therefore, abnormalities in the balance of PTZ cell fate decisions may contribute to Dandy-Walker malformation. Future studies of the maintenance and differentiation of the PTZ in mice should shed light on pathological mechanisms underlying human cerebellar birth defects.

## MATERIALS AND METHODS

### Mouse and tissue preparation

All procedures involving animals were approved by the Animal Care Committee at the University of Connecticut Health Center (protocol #101849–0621) and complied with national and state laws and policies. All mouse strains were maintained on CD1 outbred genetic background. Noon of the day on which a vaginal plug was detected was designated as E0.5 in the staging of embryos. Embryonic mouse brains were dissected in ice-cold phosphate-buffered saline and fixed in 4% paraformaldehyde between 40 minutes and overnight. Brains were cryoprotected, frozen in OCT freezing medium (Sakura Finetek), and sectioned with cryostat microtome (Leica).

Generation and characterization of the *Atoh1-Cre* (B6.Cg-Tg(Atoh1-cre)1Bfri/J*, #011104*) ^32^, *Gbx2^creER^* (*Gbx2^tm1.1(cre/ERT2)Jyhl^*/J, #022135) ^64^, *Ptf1a^CreER^* (*Ptf1a^tm2(cre/ESR1)Cvw^*/J, #019378)^81^, *R26R^RFP^* (*B6.Cg-Gt(ROSA)26Sor^tm9(CAG-tdTomato)Hze^*/J, #007909) ^33^, *R26^NICD^* (*Gt(ROSA)26Sor^tm1(Notch1)Dam^*/J, #008159) ^65^, and *Wnt1-CreER* (Tg(Wnt1-cre/ERT)1Alj/J, #008851) ^82^ alleles have been described. Primer sequences for PCR genotyping and protocols are described on the JAX Mice website.

### In situ hybridization and immunohistochemistry

Standard protocols were used for X-gal histochemistry, immunofluorescence, and *in situ* hybridization as described previously ^64^. Detailed protocols are available on the Li Laboratory website (http://lilab.uchc.edu/protocols/index.html). Primary and secondary antibodies used in the study are listed in Supplementary Table S7. Standard *in vitro* transcription methods using T7 polymerase (Roche) and digoxigenin-UTP RNA labeling mix (Roche) was used to produce antisense riboprobes. Images were collected on a Zeiss Axio Imager M1 microscope and processed using Photoshop or Fiji software.

### Cell counting

Nuclear segmentation was performed based on DAPI staining using the Fiji *Stardist* plugin ^83^. The Fiji *Annotater* plugin ^84^ was used to correct *Stardist-*produced segmentations and to identify markers associated with the segmented nuclei, manually for RFP and by using thresholding for Cre- and Meis2-stained nuclei.

### Single-cell RNA-seq and data processing

Single-cell preparation was performed as described ^10^, and Papain (Worthington Biochemical) instead of Accumax was used for tissue digestion. Sequencing libraries of E12.5 and E14.5 cerebella were generated with Chromium v2 and v3 chemistry, respectively. For Carter’s (PRJEB23051) and Vladoiu’s (GSE118068) dataset, BAM files were downloaded from European Nucleotide Archive and converted to FASTQ files using the *bamtofastq* tool (10X Genomics). RNA-seq reads were aligned to the mm10 reference genome and quantified using *cellranger count* function (CellRanger v3.1). Velocyto (v0.17.17) was used to obtain splicing-specific counts data for RNA velocity analysis. Count data were processed with the Seurat R package ^85^. From the SeuratWrapper package, *FastMNN* function, which corrects batch effects by matching mutual nearest neighbors ^86^, was used to integrate datasets. Differential gene expression analysis was performed with the *wilcoxauc* function from the presto package ^87^.

### scVelo and CellRank analyses

The dynamical model from scVelo (v0.2.2) ^28^ was used to estimate RNA velocities and velocity graphs. CellRank (v1.1.0) ^29^ was performed as described in (https://cellrank.readthedocs.io/en/latest/pancreas_advanced.html). A weighted transition matrix with 80% RNA velocity and 20% similarity was used. Clustering and Filtering of Left and Right Eigenvectors (CFLARE) estimator was used to compute fate probabilities. The terminal states were manually set based on a priori knowledge. Absorption probabilities – how likely each cell is to transition towards each terminal state were calculated. A Person’s correlation between absorption probabilities and gene expression levels was computed; genes with high correlation were considered driver genes (Supplementary Table 2). Generalized Additive Models (GAMs) were used to fit imputed gene expression trends with default settings. ClusterProfiler package ^88^ was used to examine the enrichment for biological process Gene Ontology (GO) Term and KEGG pathways of the top 100 driver genes.

### Single-nucleus ATAC-seq and data analysis

Single nuclei isolation for ATAC-seq library generation was performed following the instructions of 10X Genomics. BCL files generated from sequencing were used as inputs to the 10X Genomics Cell Ranger ATAC pipeline (version 1.2.0). FASTQ files were generated and aligned to the mm10 reference genome using BWA. The resultant fragment files were loaded into the ArchR pipeline (v1.0.0) ^89^. A cell-by-bin matrix was generated for each sample by segmenting the genome into 500-bp windows and scoring each cell for reads in each window. Cells were filtered based on log10(UMI) between 2.8 and 5.5, and the fraction of reads in promoters between 12-45%. Bins were then filtered by removing those overlapping with the ENCODE blacklist, those mapped to “random” and Y chromosomes, and the top 5% overlapping with invariant features. A cell-by-cell similarity matrix was generated by calculating the Latent Semantic Index (LSI) of the binarized bin matrix. Principal component analysis (PCA) was then performed on LSI values, cell clusters were identified with Leiden clustering.

### Defining snATAC-seq cell cluster identity

Gene activity scores were calculated using the ArchR algorithm, which primarily bases on the local accessibility of the promoter and gene body and also takes into account the distal elements (up to 5kb) and the gene size ^89^. The gene activity scores were normalized by read depth across all genes to a constant of 10,000. The *getMarkerFeatures* ArchR function was used to select marker genes based on gene activity scores with a cutoff at FDR ≤ 0.01 and log2 fold change ≥ 1.25. To match snATAC-seq and scRNA-seq data, we integrated the derived gene activity scores with gene expression levels, using canonical correction analysis to match cells from ATAC to their nearest neighbors in RNA ^85^. Using a modified *FindTransferAnchors* function of Seurat, ArchR aligned snATAC-seq and scRNA-seq data of mouse cerebella at E12.5, E13.5, and E14.5. As a result, each cell in snATAC-seq was assigned a gene expression signature which was used for downstream analyses.

### ATAC peak calling

Fragments from cells were grouped by cluster and 501-bp fixed-width peaks were called on all cluster fragments using MACS2 (https://github.com/taoliu/MACS) with the parameters ‘-- nomodel --shift -37 --ext 73 --qval 1e-2 -B -- SPMR --call-summits’. Peaks from each cluster were then combined to form a master peak set and a cell-by-peak matrix was constructed. This matrix was binarized for all downstream applications.

### Determination of differentially accessible peaks and cluster-specific peaks

The *getMarkerFeatures* ArchR function was used to identify differentially accessible peaks for each cluster with a cutoff at FDR≤0.01 and log2 fold change ≥ 1.0.

### Transcription factor motif enrichment analysis

Using motif information from CIS-BP (database 2.0), we used the *addMotifAnnotations* ArchR function to convert the peak-by-cell matrix into a motif-by-cell binary matrix based on the motif present in the peak. The *peakAnnoEnrichement* function was used to calculate enrichment for known TF motifs in cluster-specific peaks. To evaluate motif enrichments at the single-cell level based on ChromVAR ^38^, the *addDeviationsMatrix* ArchR function was used to compute per-cell motif deviations

### Identification and validation of CREs of embryonic mouse cerebellum

The ArchR *addPeak2GeneLinks* function was used to identify peak-to-gene links with the default settings except for changing the maximal distance between the peak and gene to 500 kb. After identifying 71,162 peak-to-gene links from the merged snATAC-seq dataset, we iterated the analysis and identified additional peak-to-gene links that were specific to each stage (18,158/all; 42,464/E12.5 only; 16,794/E13.5 only; 49,192/E14.5 only), resulting in a total of 197,299 unique peak-to-gene pairs.

The evolutionary conservation scores, phastCons60way and phyloP60way, were retrieved from the UCSC genome browser data portal. The average conservation scores in 2 kb centering CRE midpoint were computed using the *aggregate* function of bwtool (version 1.0)^90^. Random genomic regions were created by shuffling the original CREs. The proximal enhancer-like (72,794) and distal enhancer-like (209,040) of all mouse CREs ^42^ were downloaded from SCREEN database (https://screen.encodeproject.org). Permutation tests of overlapping between CREs and other genomic features were performed using the regionR package ^91^.

### Generation and simulation of gene regulatory networks (GRN) with CellOracle

Reconstruction and simulation of GRNs were performed with CellOracle (v0.6.2) ^50^ following the authors’ instructions (https://morris-lab.github.io/CellOracle.documentation/). Based on the inferred CRE-target list, we extracted all potential connections between TFs and targets using gimmemotifs motif database ^92^ with a false positive rate threshold of 0.02. Top GRN genes were selected and ranked using a combined score of degree centrality, betweenness centrality, and eigenvector centrality. These network scores were normalized between 0 and 1 – divided all values by the maximum of the corresponding scores. The average of the scaled scores was used to rank genes in the GRN of each cell type. Cell state transitions resulting from the perturbation of specific TFs were simulated through 5x iterations. For the simulation of the knockout of *Atoh1*, *Ptf1a*, or *Hes1*, we set the expression of the corresponding gene at 0. For the simulation of the gain of function of *Hes1*, we set the *Hes1* expression value at 3.0, which is a twofold increase of the detected *Hes1* expression. For the simulation of the *Atoh1^Ptf1a^* knock-in mutation, the detected *Ptf1a* expression was transferred to *Atoh1*. The reciprocal manipulation was applied for the simulation of the *Ptf1a^Atoh1^* knock-in mutation.

## ACKNOWLEDGEMENTS

We thank Ms. Jie Zhou and Dr. Alan Liang for their contribution to the study. Dr. Rutesh Vyas for critical reading and comments on the manuscript.

## Author contributions

Conception and supervision of the study: JL

Data generation: NF, QG, KM, JS, and JL

Data analysis: NF, QG, KM, and JL

Data interpretation: NF, QG, and JL Writing the manuscript: JL

Review and editing of the manuscript: NF, QG, and JL

## Data and code Availability

Sequencing data have been deposited in GEO under accession codes GSE178546.

To make the data easily accessible, we have generated searchable html pages where users can explore and visualize the scRNA-seq E10.5-E17.5 and E10.5-E13.5. Putative CREs and CRE-target links across cell types can be inspected as a custom UCSC genome browser tracks. Computer code associated with the manuscript are available on the LiLab GitHub.

---

**Supplementary Table 1**: Lists of molecular features of different cell types and states of embryonic mouse cerebella (E10.5-E17.5/sheet1 and E10.5-E13.5/sheet2).

**Supplementary Table 2**: List of driver genes identified by CellRank.

**Supplementary Table 3**: Lists of cluster-specific markers based on gene activity scores (sheet = sheet = GeneScoreMatrix) and RNA-integrated expression (sheet = GeneIntegratedMatrix) in snATAC-seq.

**Supplementary Table 4**: List of putative transcription activators and repressors in early cerebellar development.

**Supplementary Table 5**: List of top 10 percentile genes with the highest number of CREs and results of GO term enrichment analysis.

**Supplementary Table 6**: List of CellOracle-inferred key regulators of different cell groups.

**Supplementary Table 7**: List of antibodies used in this study.

## SUPPLEMENTARY MATERIALS

**Supplementary Figure S1.**
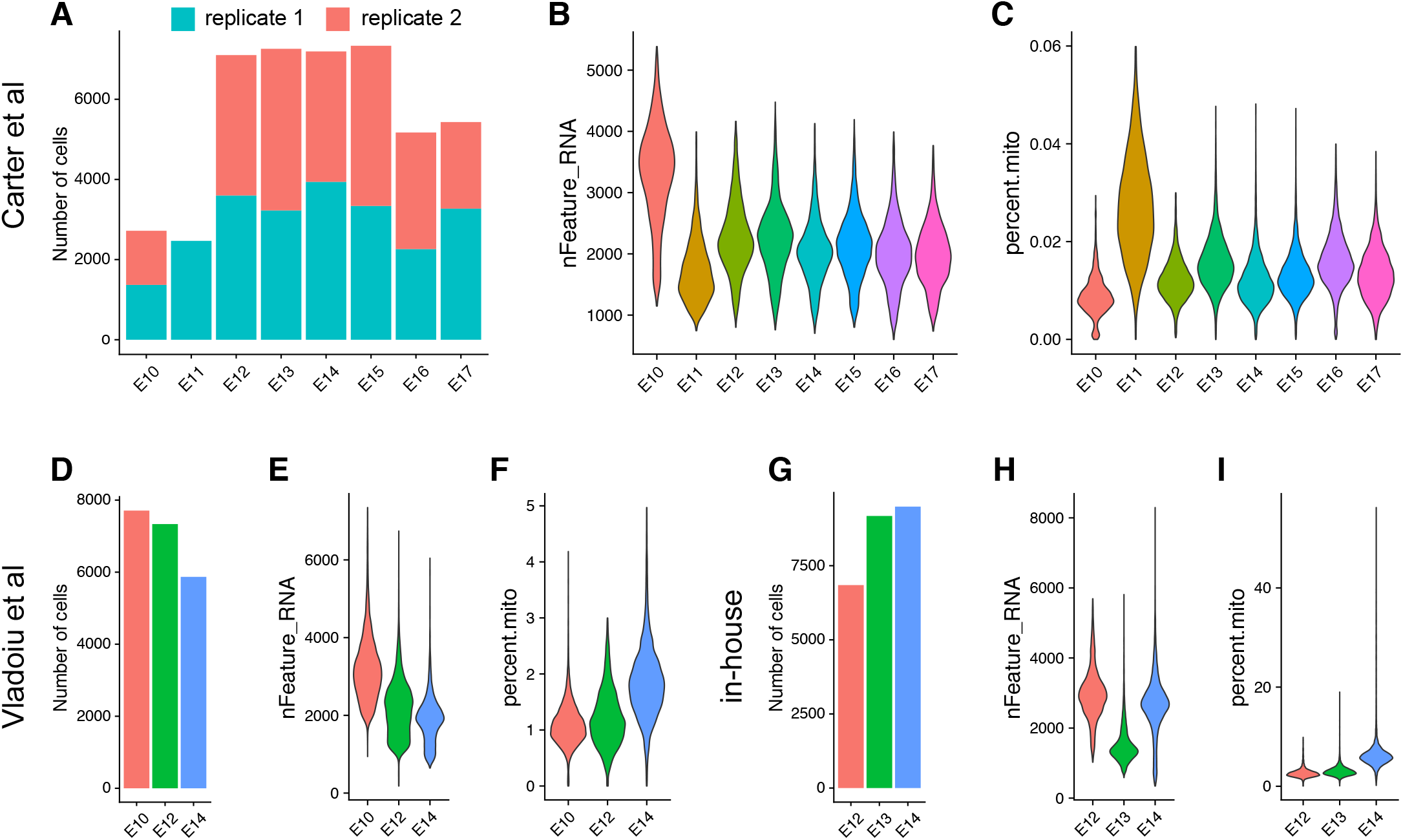
Comparison of the data quality among three scRNA-seq datasets used in the current study. Barplot and violin plots showing the number (A) and quality (B,C) of scRNA-seq data from Carter et al. Barplot and violin plots showing the number (D) and quality (E,F) scRNA-seq from Vladoiu et al. Barplot and violin plots showing the number (G) and quality (H,I) of scRNA-seq generated in-house.

**Supplementary Figure S2.**
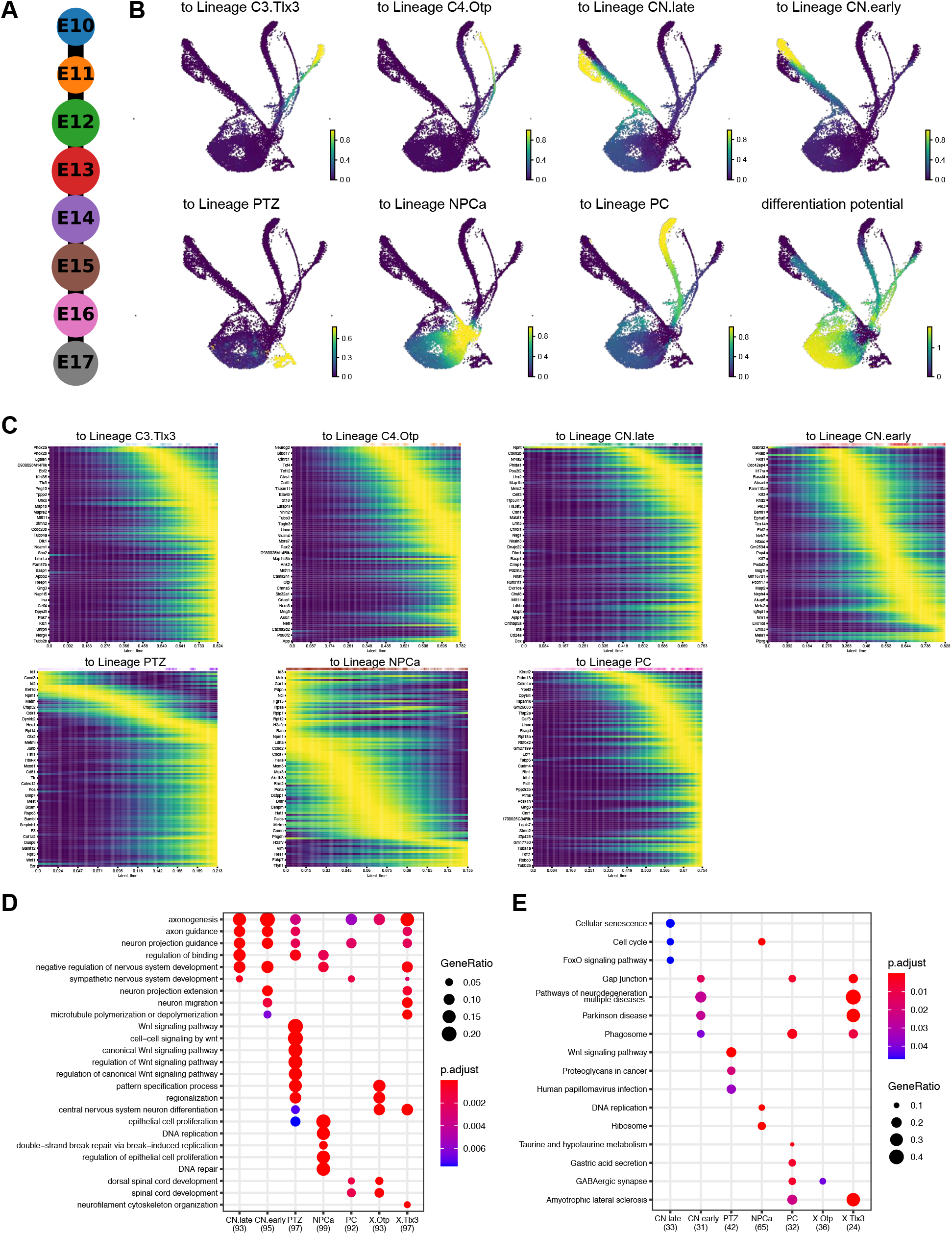
Inference of cerebellar development with scRNA-seq. (A) Unbiased hierarchical ordering of cerebellar cells from different embryonic stages by PAGA. (B) Cell-fate probabilities inferred by CellRank in different lineages. (C) Heatmaps showing the expression dynamics of top 100 driver genes in different lineages. Dot plots showing enrichment for GO terms (D) and KEGG pathways (E) of top 100 driver genes.

**Supplementary Figure S3.**
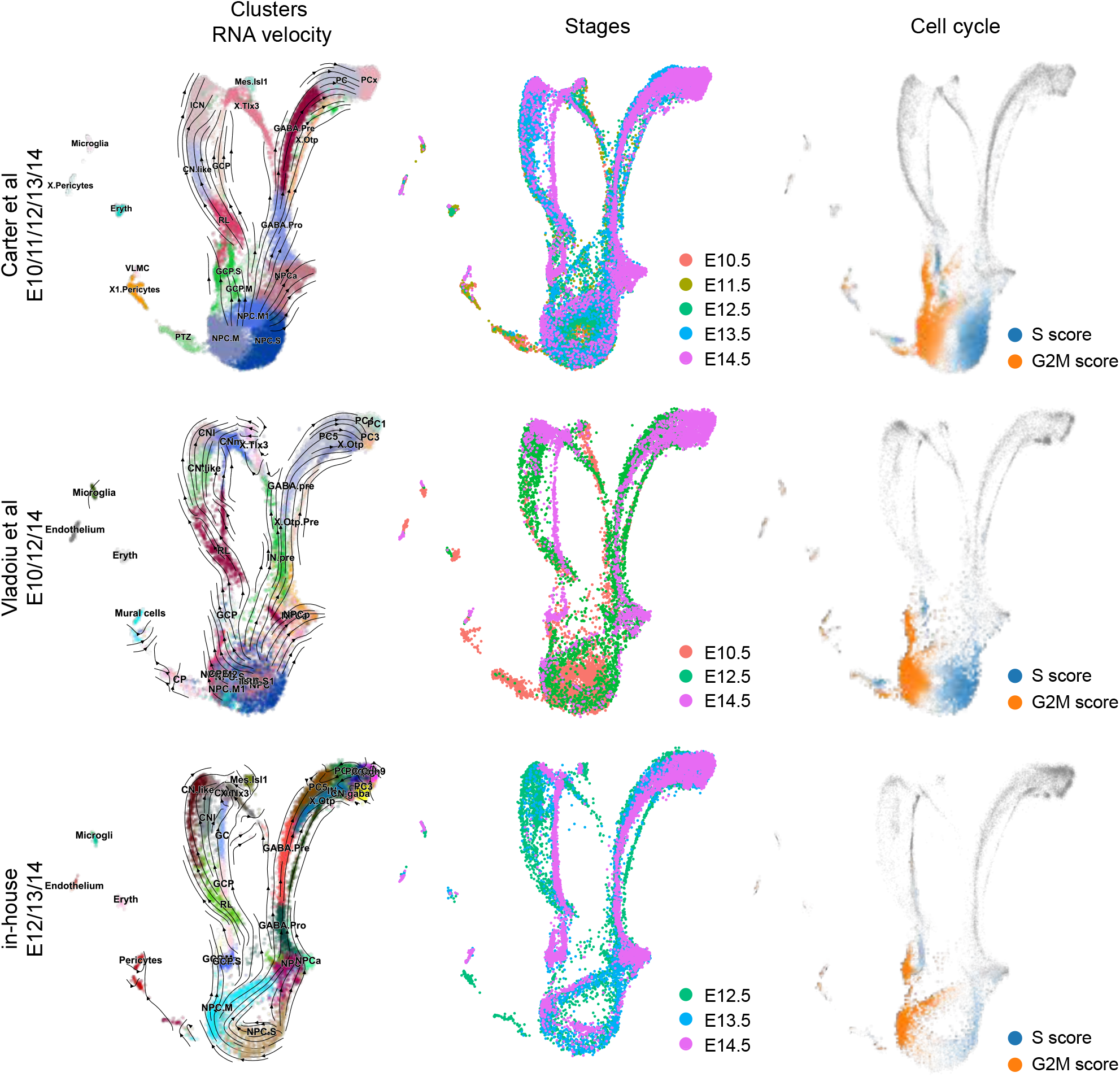
Assessment of the robustness of scRNA-seq analyses. UMAP showing cell clusters, RNA velocity streams, stages, and cell cycles of three different scRNA-seq datasets.

**Supplementary Figure S4.**
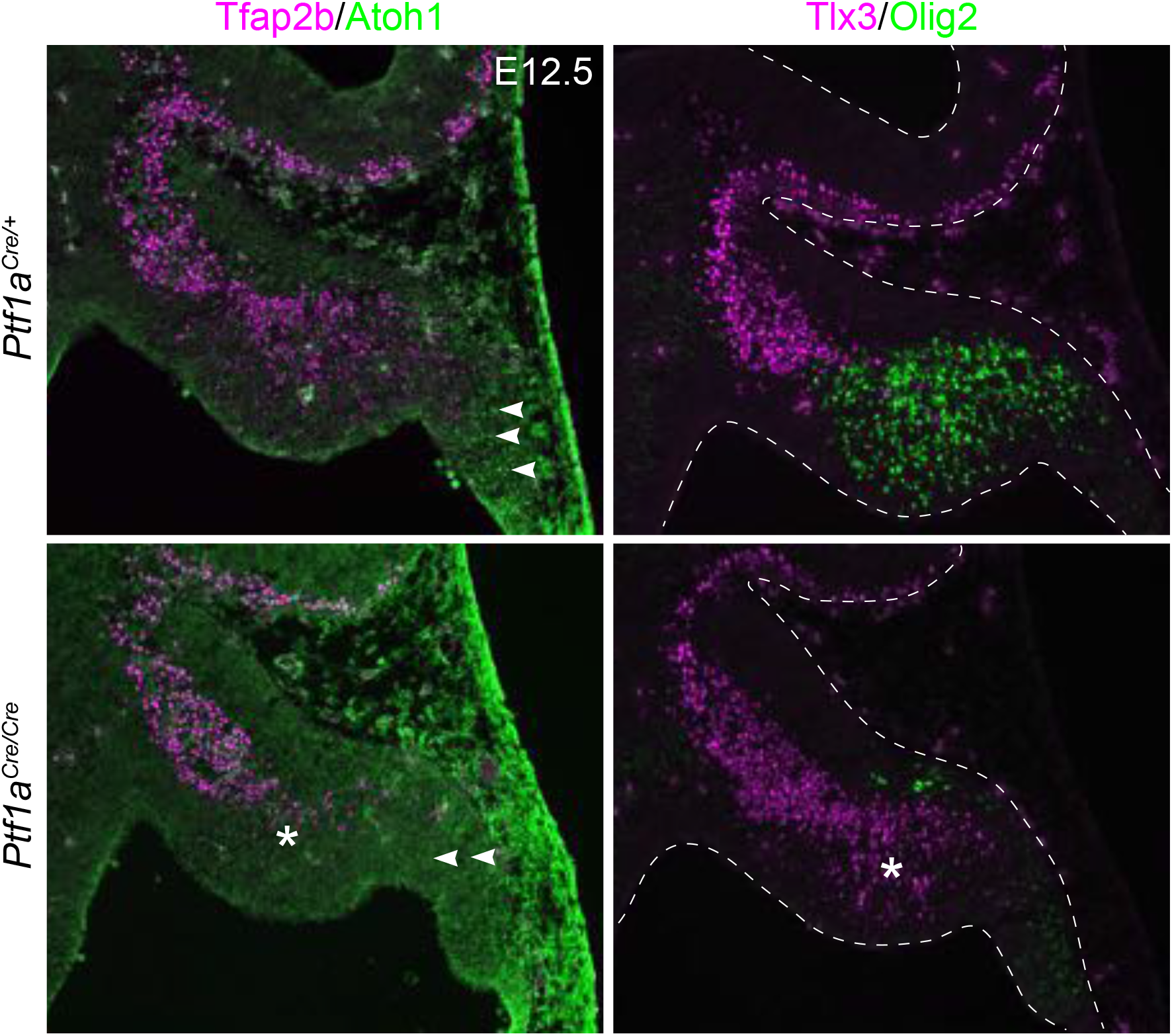
Conversion of the C2 fate to C3 in the cerebellar anlage without *Ptf1a*. Immunofluorescence on sagittal sections of E12.5 cerebella of the indicated genotype. Arrowheads indicate nascent Atoh1-expressing cells; the asterisk denotes the reduction of Tfap2b and the absence of Olig2 in the C2 area.

**Supplementary Figure S5.**
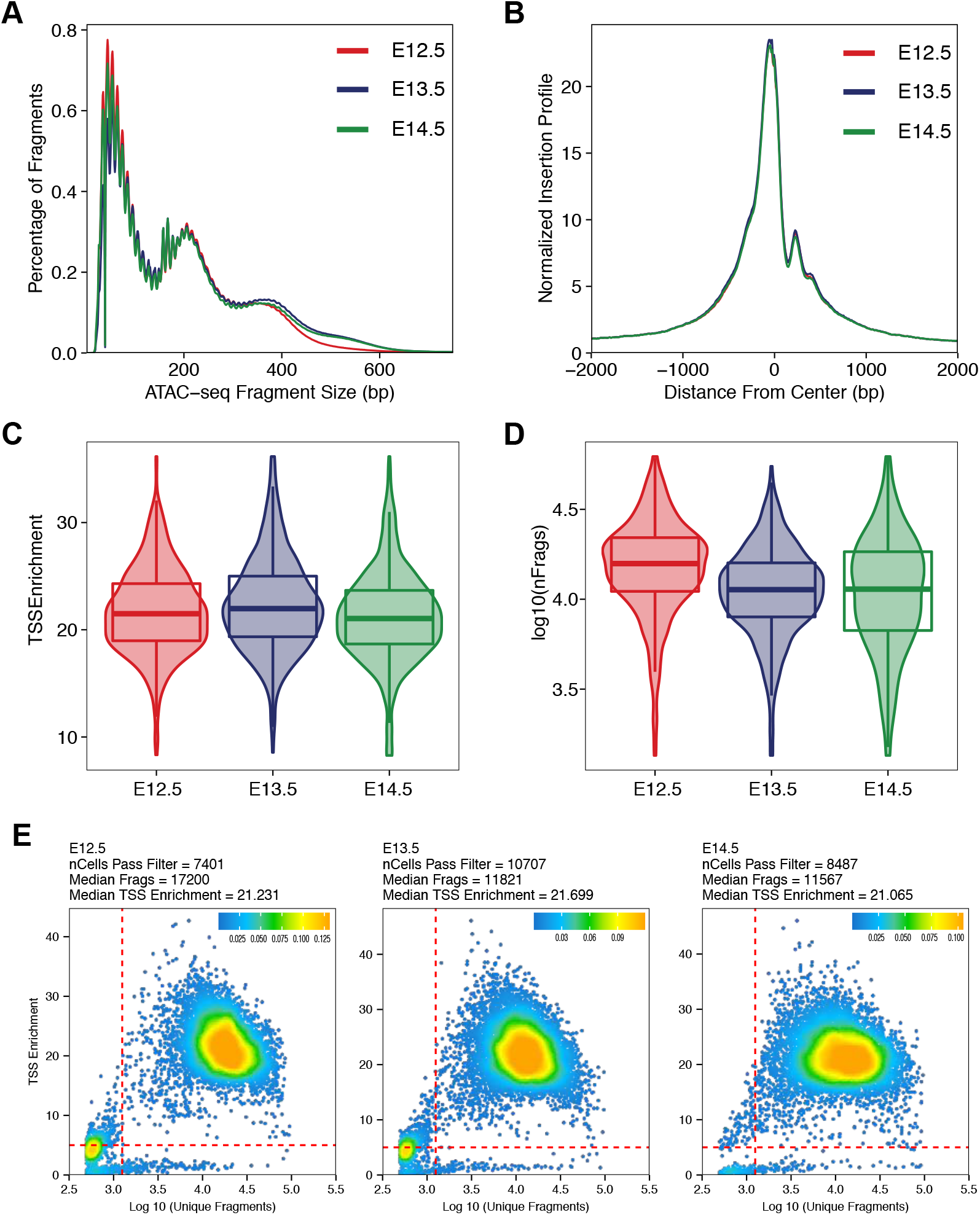
Data quality of the snATAC-seq libraries. Line plots showing the fragment size distribution (A) and fragment distribution around the transcription start site (TSS; B). Boxplots comparing the TSS enrichment (C) and total fragments (D) of different experiments. (E) Scatter plots showing filtering of snATAC-seq cells based on TSS enrichments and total fragments. The red dashed lines indicate the filtering threshold.

**Supplementary Figure S6.**
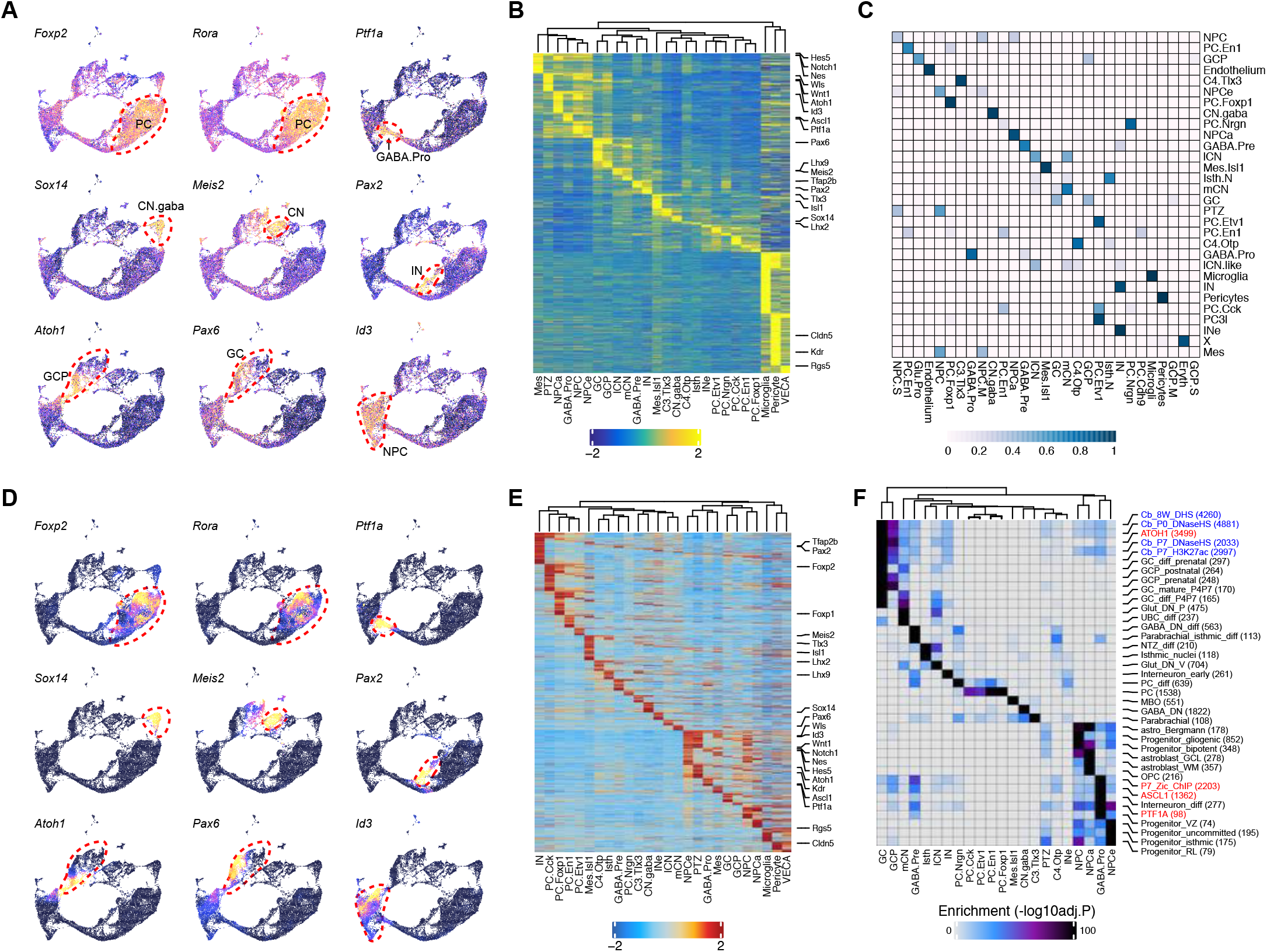
Linkage of ATAC cells with RNA cells. UMAP (A) and heatmap (B) showing marker expression based on gene activity scores. Each cerebellar cell type is circled by red dash lines. (C) Heatmap showing the alignment between scATAC-seq and scRNA-seq clusters. UMAP (D) and heatmap (E) showing linked gene expression calculated based on integrated snATAC-seq and scRNA-seq. Note the similar pattern between A and D, whereas the latter displays more dynamic differences between cell clusters. (F) Heatmap showing enrichments of cell-specific peaks with various genomic features published previously. Published ChIP-seq data are in red; bulk DNAse I hypersensitive site-seq from ENCODE in blue; cell-specific peaks from a previous snATAC-seq in black.

**Supplementary Figure S7.**
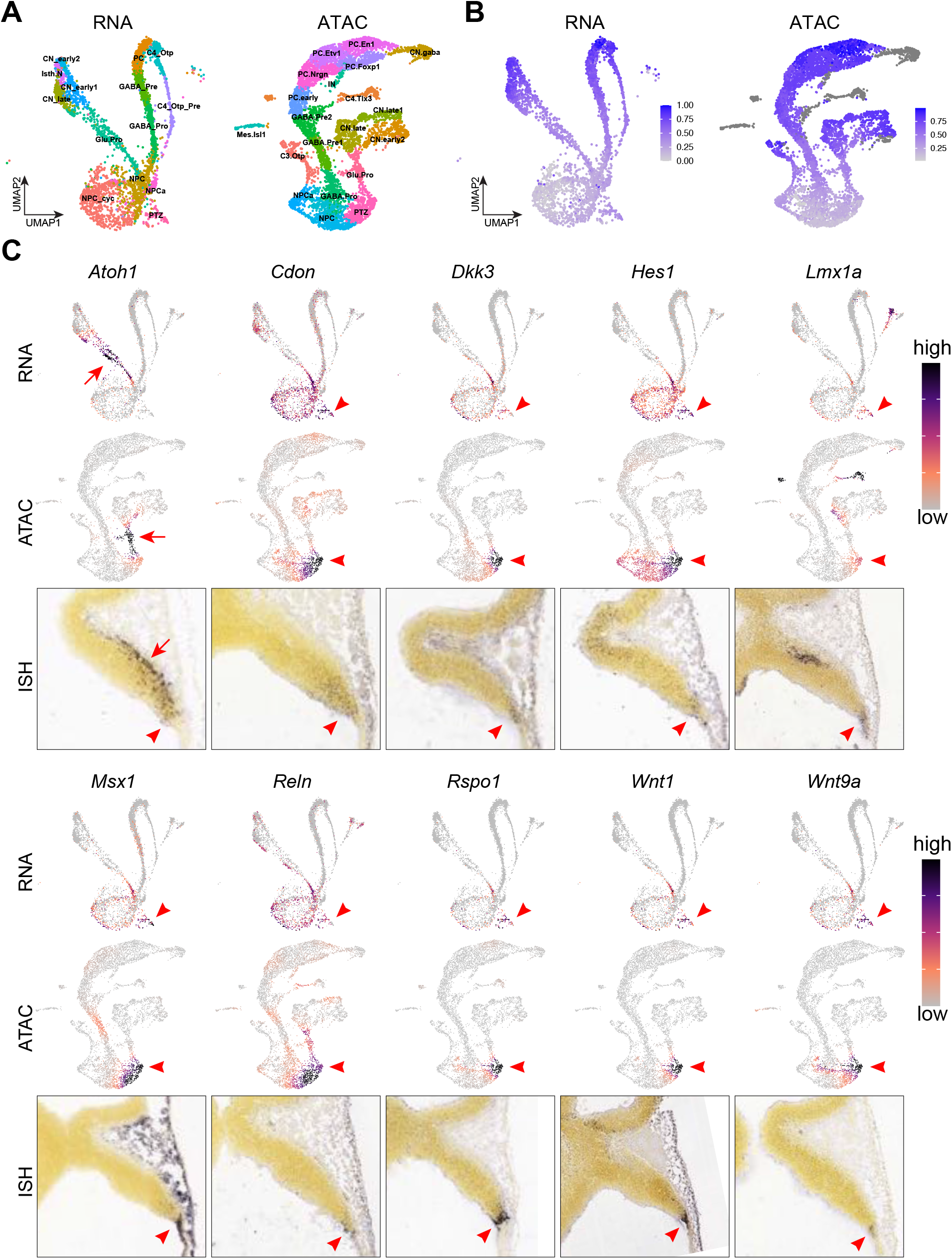
Characterization of the molecular feature of PTZ cells with both scRNA-seq and snATAC-seq. UMAP embeddings of scRNA and snATAC cells from E12.5 mouse cerebella exhibit similar global structure (A) and developmental continuum as measured by latent time inferred by scVelo (B). Inspection of feature genes of the PTZ domain in scRNA, snATAC cells and, *in situ* hybridization on sagittal sections of E11.5 cerebella (from Allen Developing Mouse Brain Atlas). Arrowheads indicate the PTZ; arrows show glutamatergic progenitors.

**Supplementary Figure S8.**
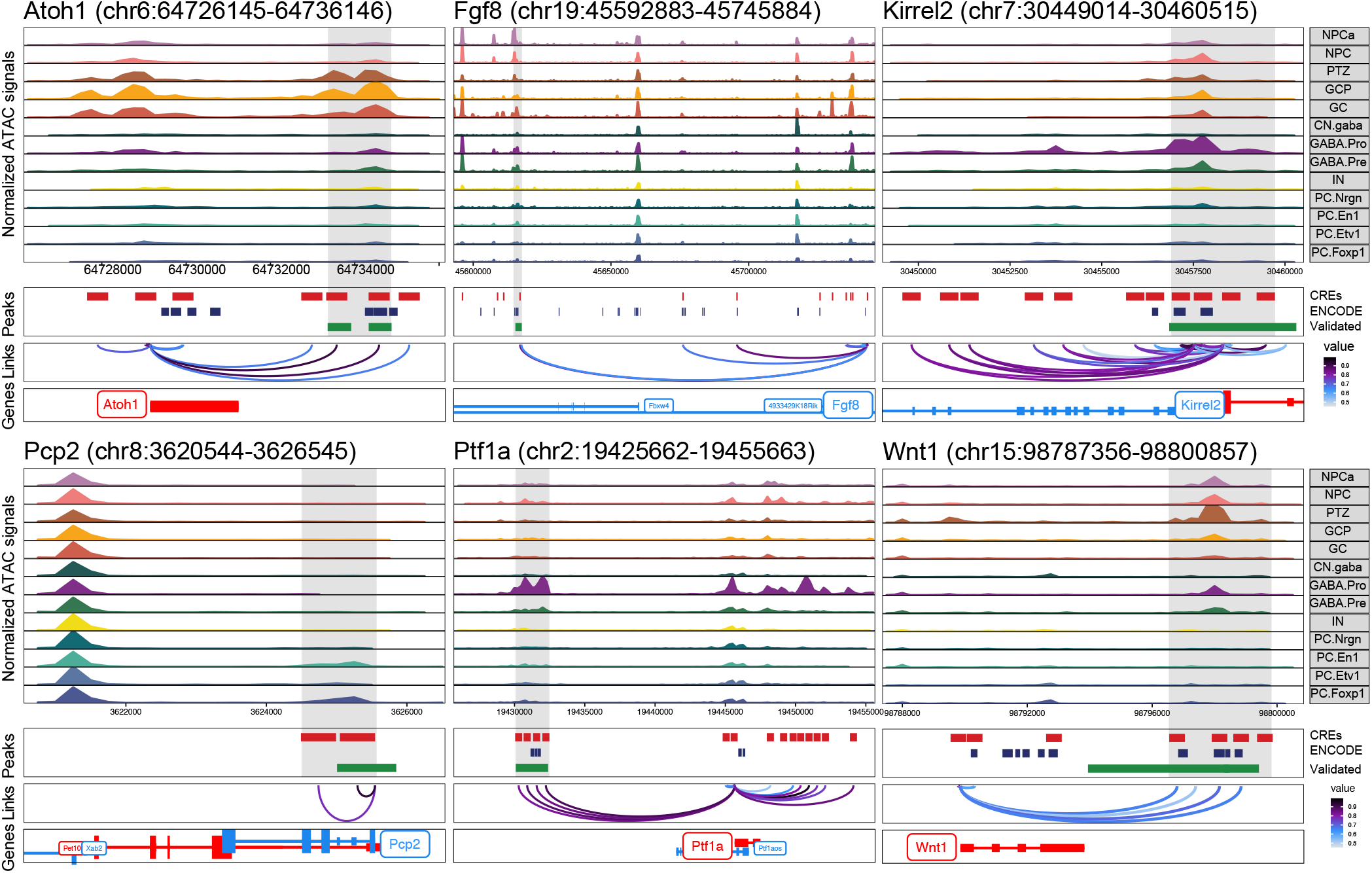
Overlapping between CREs and validated enhancers. Examples of linkages of CREs and targets. The grey areas highlight the CRE that has been validated by transgenic mouse assays.

**Supplementary Figure S9.**
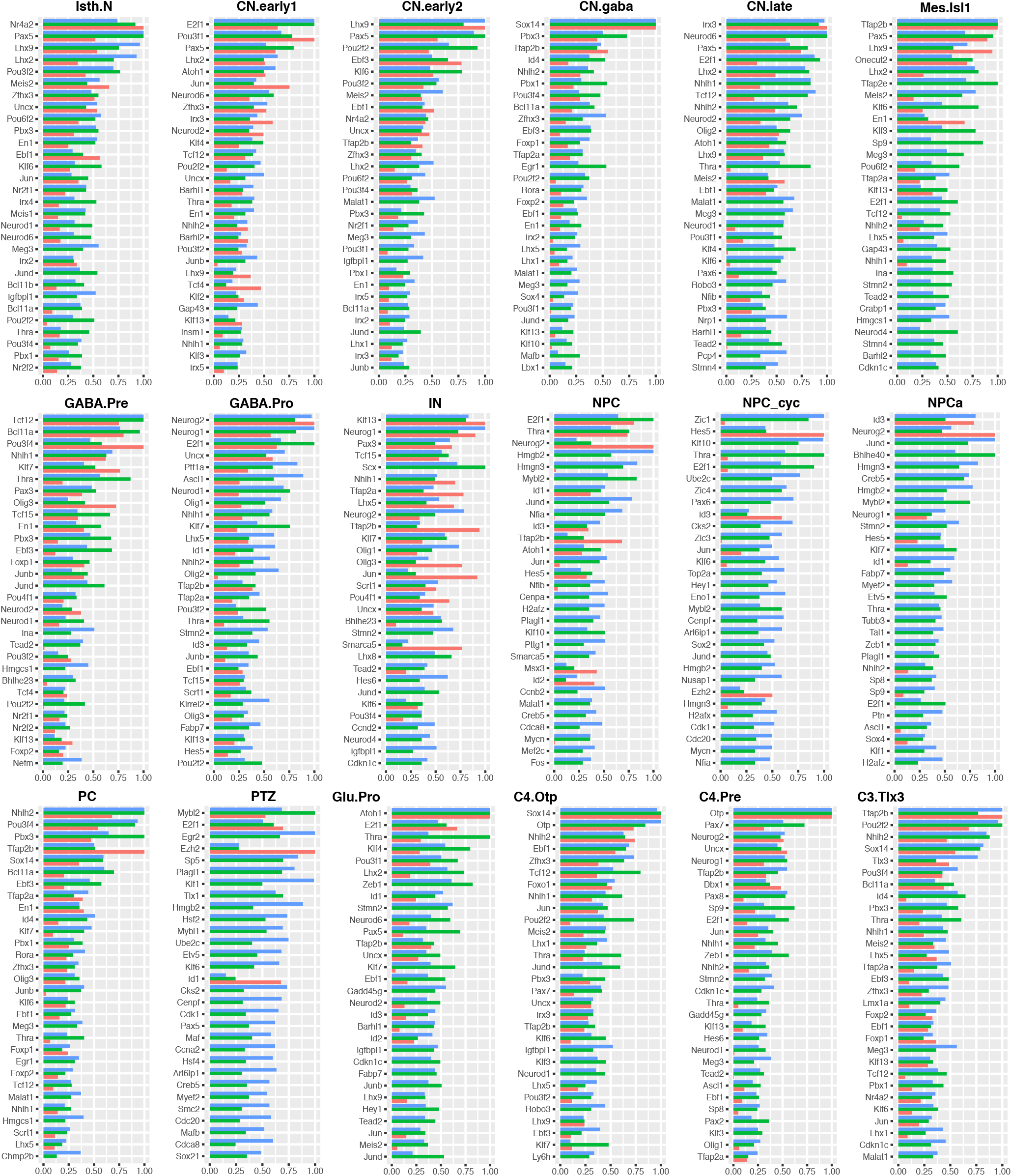
Identification of putative key regulators through network analysis of the GRN specific to each cell type or state. Top 30 genes based on the aggregate ranking of three key network scores: degree centrality, betweenness centrality, and eigenvector centrality, in the GRN for each cell cluster.

**Supplementary Figure S10.**
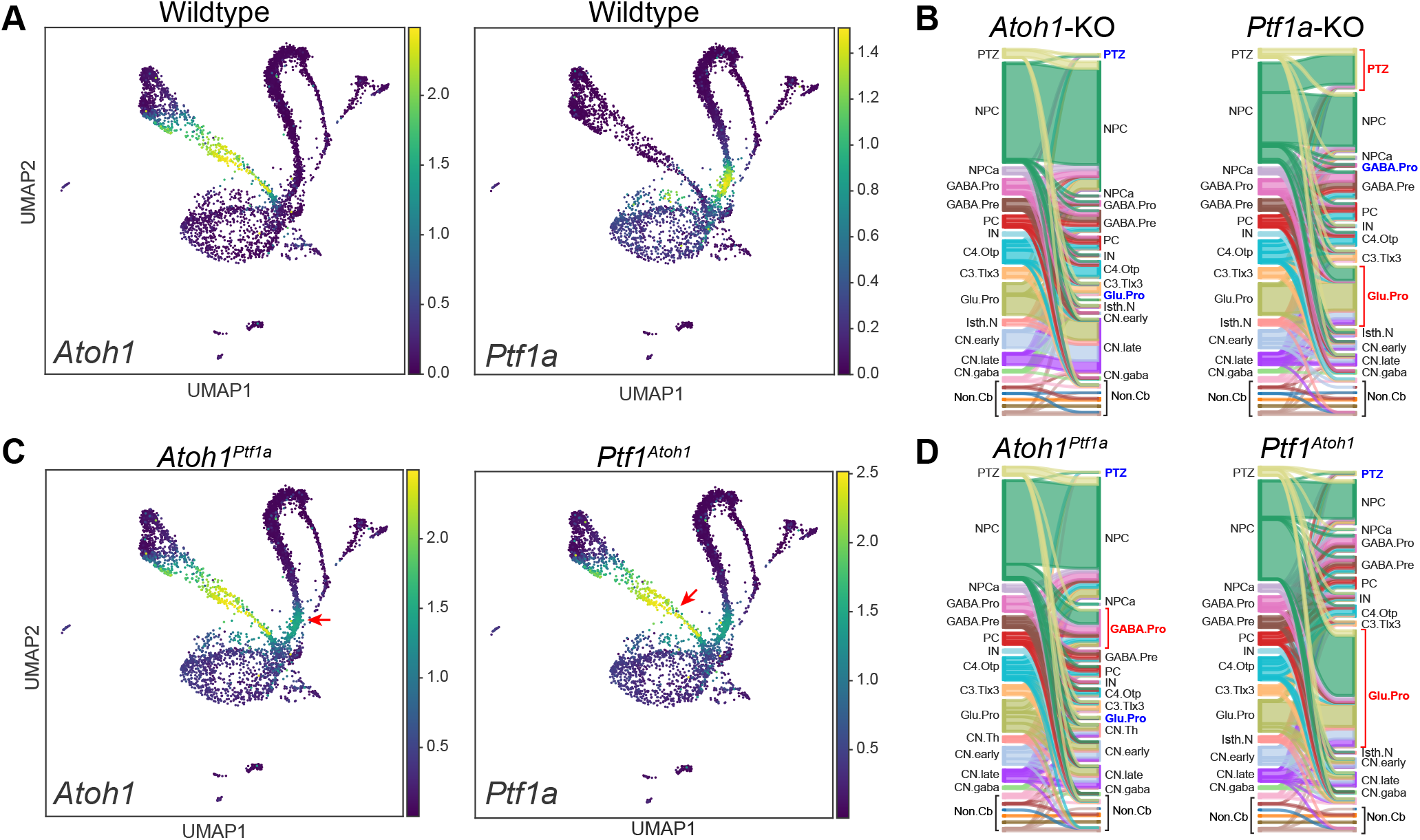
Simulation of genetic perturbations of *Atoh1* and *Ptf1a* during cerebellar development. (A) UMAP showing the expression of *Atoh1* and *Ptf1a* in E12.5 scRNA-seq. (B) Sankey diagrams showing simulated cell transition between different cell types caused by genetic perturbations. Simulated expression of *Atoh1* and *Ptf1a* (C) and cell-fate changes (D) in the cerebellar carrying knock-in alleles *Ptf1a^Atoh1^* and *Atoh1^Ptf1a^*. Expanded or reduced cell types are shown in red and blue, respectively.

**Supplementary Figure S11.**
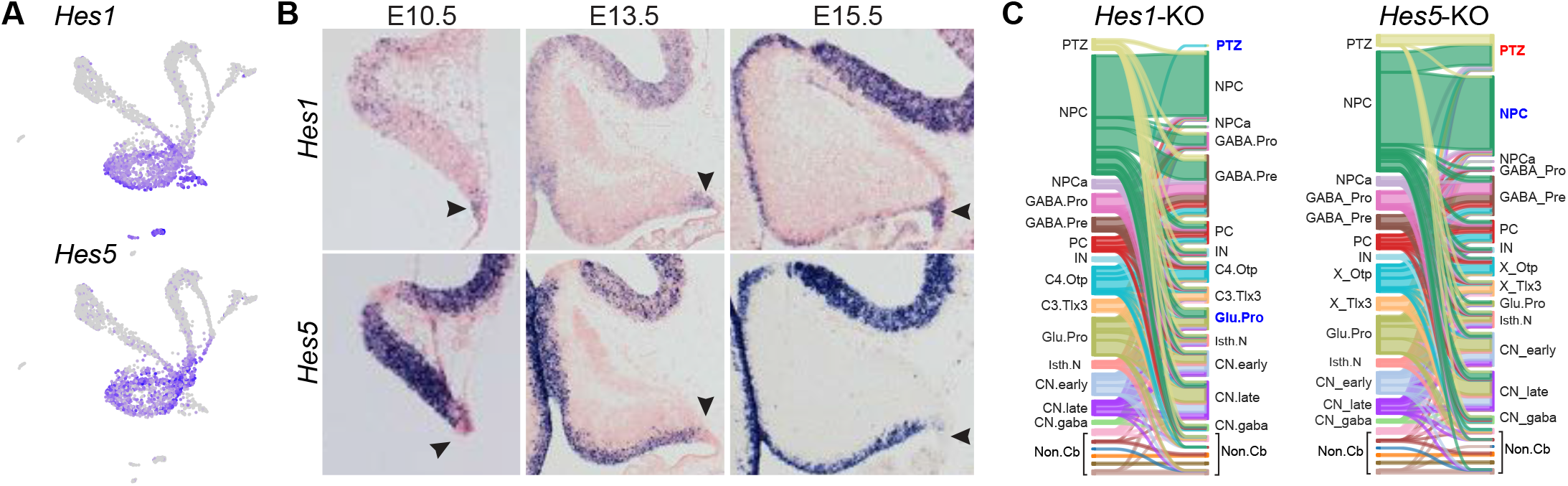
Important role of *Hes1* in PTZ development. Expression of *Hes1*, but not *Hes5*, in the PTZ as shown by scRNA-seq of E12.5 cerebella (A) and in situ hybridization (B). Sankey diagrams showing simulated cell transition between different cell types caused by the loss of *Hes1* or *Hes5* (C). Note the opposing effect on the PTZ caused by the deletion of *Hes1* and *Hes5*.

**Supplementary Figure S12.**
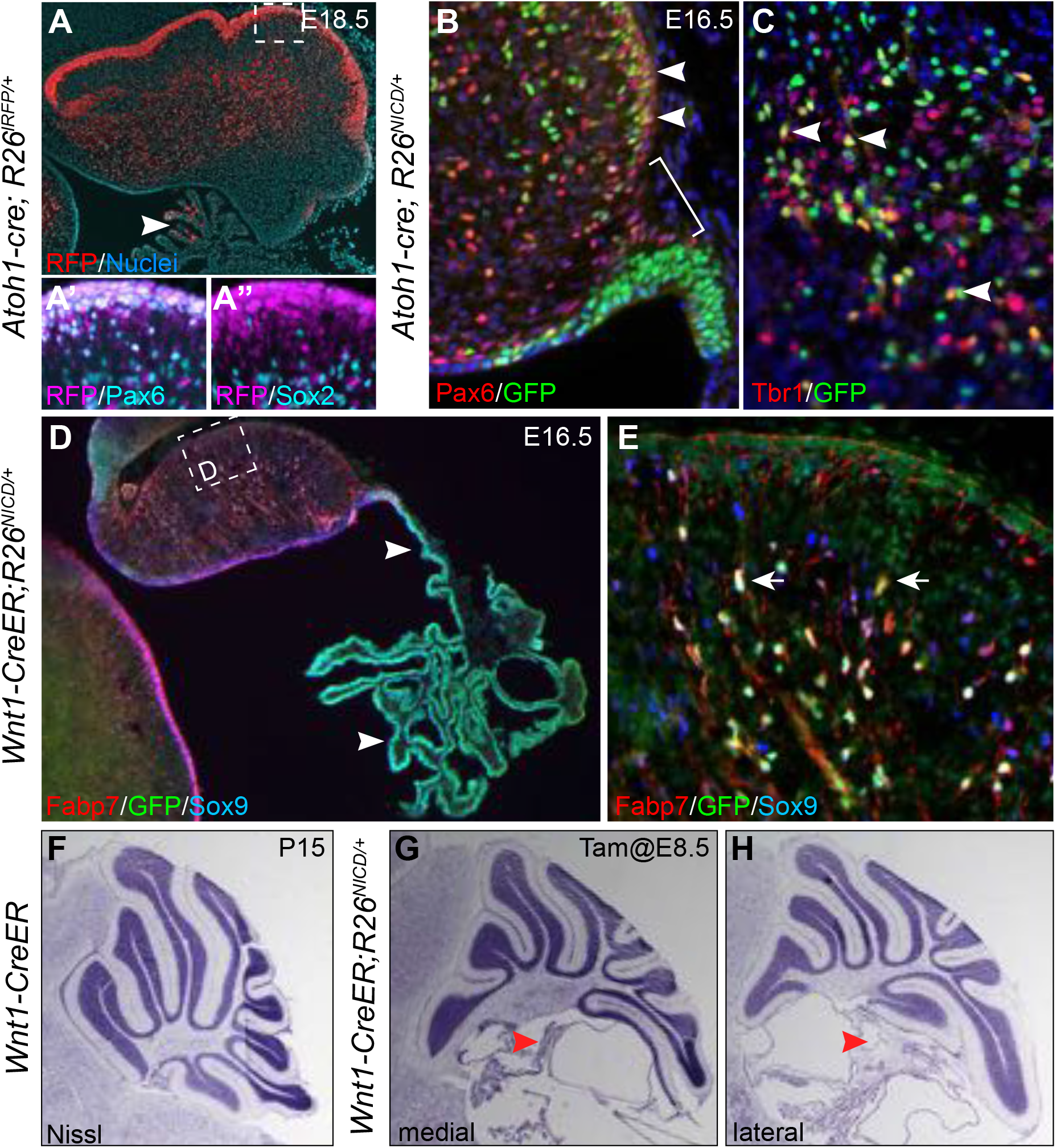
NICD gain of function analysis. (**A-A”**) Red fluorescence and immunofluorescence showing the distribution of *Atoh1*-lineage descendants in the cerebellum. (**B and C**) Abundant *NICD*^+^ cells are present in the EGL and the fastigial nucleus of *Atoh1-Cre; R26^NICD/+^* embryos at E16.5. The bracket indicates the absence of *NICD*^+^ cells in the EGL next to the PTZ. (**D and E**) Immunofluorescence on a sagittal section of E16.5 *Wnt1-CreER;R26^NICD/+^* embryos that were given tamoxifen at E8.5. The arrowhead indicates the enlarged choroid plexus; arrows denote Bergmann glia with NICD expression. (**F-H**) Nissl histology of sagittal sections of the cerebellar vermis of indicated genotypes at P15. The arrowhead shows the enlarged choroid plexus. Note that the posterior truncation is more prominent in the medial section (G) than the lateral (H).

## REFERENCES

1. Ito, M. Control of mental activities by internal models in the cerebellum. Nature Reviews Neuroscience 9, 304–313 (2008).

2. Reeber, S. L., Otis, T. S. & Sillitoe, R. V. New roles for the cerebellum in health and disease. Front. Syst. Neurosci. 7, (2013).

3. Leto, K. et al. Consensus Paper: Cerebellar Development. Cerebellum 15, 789–828 (2016).

4. Englund, C. et al. Unipolar Brush Cells of the Cerebellum Are Produced in the Rhombic Lip and Migrate through Developing White Matter. J. Neurosci. 26, 9184–9195 (2006).

5. Machold, R. & Fishell, G. Math1 Is Expressed in Temporally Discrete Pools of Cerebellar Rhombic-Lip Neural Progenitors. Neuron 48, 17–24 (2005).

6. Ben-Arie, N. et al. Math1 is essential for genesis of cerebellar granule neurons. Nature 390, 169–172 (1997).

7. Hoshino, M. et al. Ptf1a, a bHLH Transcriptional Gene, Defines GABAergic Neuronal Fates in Cerebellum. Neuron 47, 201–213 (2005).

8. Wang, B., Lin, D., Li, C. & Tucker, P. Multiple Domains Define the Expression and Regulatory Properties of Foxp1 Forkhead Transcriptional Repressors. J. Biol. Chem. 278, 24259–24268 (2003).

9. Yamada, M. et al. Specification of Spatial Identities of Cerebellar Neuron Progenitors by Ptf1a and Atoh1 for Proper Production of GABAergic and Glutamatergic Neurons. J. Neurosci. 34, 4786–4800 (2014).

10. Wizeman, J. W., Guo, Q., Wilion, E. M. & Li, J. Y. Specification of diverse cell types during early neurogenesis of the mouse cerebellum. eLife 8, e42388 (2019).

11. Haldipur, P. et al. Spatiotemporal expansion of primary progenitor zones in the developing human cerebellum. Science 366, 454–460 (2019).

12. Waddington, C. H. The strategy of the genes; a discussion of some aspects of theoretical biology. (Allen & Unwin, 1957).

13. Buenrostro, J. D. et al. Single-cell chromatin accessibility reveals principles of regulatory variation. Nature 523, 486–490 (2015).

14. Briggs, J. A. et al. The dynamics of gene expression in vertebrate embryogenesis at single-cell resolution. Science 360, (2018).

15. Farrell, J. A. et al. Single-cell reconstruction of developmental trajectories during zebrafish embryogenesis. Science 360, (2018).

16. Wagner, D. E. et al. Single-cell mapping of gene expression landscapes and lineage in the zebrafish embryo. Science 360, 981–987 (2018).

17. Carter, R. A. et al. A Single-Cell Transcriptional Atlas of the Developing Murine Cerebellum. Current Biology 28, 2910–2920.e2 (2018).

18. Kozareva, V. et al. A transcriptomic atlas of the mouse cerebellum reveals regional specializations and novel cell types. bioRxiv 2020.03.04.976407 (2020) doi:10.1101/2020.03.04.976407.

19. Peng, J. et al. Single-cell transcriptomes reveal molecular specializations of neuronal cell types in the developing cerebellum. Journal of Molecular Cell Biology 11, 636–648 (2019).

20. Sarropoulos, I. et al. Developmental and evolutionary dynamics of cis-regulatory elements in mouse cerebellar cells. Science (2021) doi:10.1126/science.abg4696.

21. Vladoiu, M. C. et al. Childhood cerebellar tumours mirror conserved fetal transcriptional programs. Nature 572, 67–73 (2019).

22. Aldinger, K. A. et al. Spatial and cell type transcriptional landscape of human cerebellar development. Nat Neurosci 1–13 (2021) doi:10.1038/s41593-021-00872-y.

23. Wolf, F. A. et al. PAGA: graph abstraction reconciles clustering with trajectory inference through a topology preserving map of single cells. Genome Biology 20, 59 (2019).

24. Chizhikov, V. V. et al. The roof plate regulates cerebellar cell-type specification and proliferation. Development 133, 2793–2804 (2006).

25. Zordan, P., Croci, L., Hawkes, R. & Consalez, G. G. Comparative analysis of proneural gene expression in the embryonic cerebellum. Developmental Dynamics 237, 1726–1735 (2008).

26. Dai, J.-X., Hu, Z.-L., Shi, M., Guo, C. & Ding, Y.-Q. Postnatal ontogeny of the transcription factor Lmx1b in the mouse central nervous system. J Comp Neurol 509, 341–355 (2008).

27. Millen, K. J., Steshina, E. Y., Iskusnykh, I. Y. & Chizhikov, V. V. Transformation of the cerebellum into more ventral brainstem fates causes cerebellar agenesis in the absence of Ptf1a function. PNAS 111, E1777–E1786 (2014).

28. Bergen, V., Lange, M., Peidli, S., Wolf, F. A. & Theis, F. J. Generalizing RNA velocity to transient cell states through dynamical modeling. Nature Biotechnology 38, 1408–1414 (2020).

29. Lange, M. et al. CellRank for directed single-cell fate mapping. bioRxiv 2020.10.19.345983 (2020) doi:10.1101/2020.10.19.345983.

30. Prekop, H.-T. et al. Sox14 Is Required for a Specific Subset of Cerebello–Olivary Projections. J. Neurosci. 38, 9539–9550 (2018).

31. Wang, V. Y., Rose, M. F. & Zoghbi, H. Y. Math1 Expression Redefines the Rhombic Lip Derivatives and Reveals Novel Lineages within the Brainstem and Cerebellum. Neuron 48, 31–43 (2005).

32. Matei, V. et al. Smaller inner ear sensory epithelia in Neurog1 null mice are related to earlier hair cell cycle exit. Developmental Dynamics 234, 633–650 (2005).

33. Madisen, L. et al. A robust and high-throughput Cre reporting and characterization system for the whole mouse brain. Nature Neuroscience 13, 133–140 (2010).

34. Fink, A. J. et al. Development of the Deep Cerebellar Nuclei: Transcription Factors and Cell Migration from the Rhombic Lip. J. Neurosci. 26, 3066–3076 (2006).

35. Ma, S. et al. Chromatin Potential Identified by Shared Single-Cell Profiling of RNA and Chromatin. Cell 183, 1103–1116.e20 (2020).

36. Borromeo, M. D. et al. A transcription factor network specifying inhibitory versus excitatory neurons in the dorsal spinal cord. Development 141, 2803–2812 (2014).

37. Klisch, T. J. et al. In vivo Atoh1 targetome reveals how a proneural transcription factor regulates cerebellar development. PNAS 108, 3288–3293 (2011).

38. Schep, A. N., Wu, B., Buenrostro, J. D. & Greenleaf, W. J. chromVAR: inferring transcription-factor-associated accessibility from single-cell epigenomic data. Nat Methods 14, 975–978 (2017).

39. Domcke, S. et al. A human cell atlas of fetal chromatin accessibility. Science 370, eaba7612 (2020).

40. Iwafuchi-Doi, M. & Zaret, K. S. Cell fate control by pioneer transcription factors. Development 143, 1833–1837 (2016).

41. Andersson, R. et al. An atlas of active enhancers across human cell types and tissues. Nature 507, 455–461 (2014).

42. Moore, J. E. et al. Expanded encyclopaedias of DNA elements in the human and mouse genomes. Nature 583, 699–710 (2020).

43. Echelard, Y., Vassileva, G. & McMahon, A. P. Cis-acting regulatory sequences governing Wnt-1 expression in the developing mouse CNS. Development 120, 2213–2224 (1994).

44. Helms, A. W., Abney, A. L., Ben-Arie, N., Zoghbi, H. Y. & Johnson, J. E. Autoregulation and multiple enhancers control Math1 expression in the developing nervous system. Development 127, 1185–1196 (2000).

45. Hörnblad, A., Bastide, S., Langenfeld, K., Langa, F. & Spitz, F. Dissection of the Fgf8 regulatory landscape by in vivo CRISPR-editing reveals extensive intra- and inter-enhancer redundancy. Nature Communications 12, 439 (2021).

46. Meredith, D. M., Masui, T., Swift, G. H., MacDonald, R. J. & Johnson, J. E. Multiple Transcriptional Mechanisms Control Ptf1a Levels during Neural Development Including Autoregulation by the PTF1-J Complex. Journal of Neuroscience 29, 11139–11148 (2009).

47. Mizuhara, E. et al. Purkinje cells originate from cerebellar ventricular zone progenitors positive for Neph3 and E-cadherin. Developmental Biology 338, 202–214 (2010).

48. Nitta, K., Matsuzaki, Y., Konno, A. & Hirai, H. Minimal Purkinje Cell-Specific PCP2/L7 Promoter Virally Available for Rodents and Non-human Primates. Molecular Therapy -Methods & Clinical Development 6, 159–170 (2017).

49. Pfeffer, P. L., Payer, B., Reim, G., Magliano, M. P. di & Busslinger, M. The activation and maintenance of Pax2 expression at the mid-hindbrain boundary is controlled by separate enhancers. Development 129, 307–318 (2002).

50. Kamimoto, K., Hoffmann, C. M. & Morris, S. A. CellOracle: Dissecting cell identity via network inference and in silico gene perturbation. bioRxiv 2020.02.17.947416 (2020) doi:10.1101/2020.02.17.947416.

51. Florio, M. et al. Neurogenin 2 regulates progenitor cell-cycle progression and Purkinje cell dendritogenesis in cerebellar development. Development 139, 2308–2320 (2012).

52. Lowenstein, E. D. et al. Olig3 regulates early cerebellar development. eLife 10, e64684 (2021).

53. Seto, Y. et al. Temporal identity transition from Purkinje cell progenitors to GABAergic interneuron progenitors in the cerebellum. Nature Communications 5, 3337 (2014).

54. Sudarov, A. et al. Ascl1 Genetics Reveals Insights into Cerebellum Local Circuit Assembly. J. Neurosci. 31, 11055–11069 (2011).

55. Zainolabidin, N., Kamath, S. P., Thanawalla, A. R. & Chen, A. I. Distinct Activities of Tfap2A and Tfap2B in the Specification of GABAergic Interneurons in the Developing Cerebellum. Front. Mol. Neurosci. 10, (2017).

56. Zhao, Y. et al. LIM-homeodomain proteins Lhx1 and Lhx5, and their cofactor Ldb1, control Purkinje cell differentiation in the developing cerebellum. PNAS 104, 13182–13186 (2007).

57. Green, M. J. & Wingate, R. J. Developmental origins of diversity in cerebellar output nuclei. Neural Development 9, 1 (2014).

58. Li, S., Qiu, F., Xu, A., Price, S. M. & Xiang, M. Barhl1 Regulates Migration and Survival of Cerebellar Granule Cells by Controlling Expression of the Neurotrophin-3 Gene. J. Neurosci. 24, 3104–3114 (2004).

59. Miyata, T., Maeda, T. & Lee, J. E. NeuroD is required for differentiation of the granule cells in the cerebellum and hippocampus. Genes Dev 13, 1647–1652 (1999).

60. Olson, J. M. et al. NeuroD2 Is Necessary for Development and Survival of Central Nervous System Neurons. Developmental Biology 234, 174–187 (2001).

61. . Nakamura, Y., et al. The bHLH Gene Hes1 as a Repressor of the Neuronal Commitment of CNS Stem Cells. J. Neurosci. 20, 283–293 (2000).

62. Ohtsuka, T. et al. Hes1 and Hes5 as Notch effectors in mammalian neuronal differentiation. The EMBO Journal 18, 2196–2207 (1999).

63. Kageyama, R. & Ohtsuka, T. The Notch-Hes pathway in mammalian neural development. Cell Research 9, 179–188 (1999).

64. Chen, L., Guo, Q. & Li, J. Y. H. Transcription factor Gbx2 acts cell-nonautonomously to regulate the formation of lineage-restriction boundaries of the thalamus. Development 136, 1317–1326 (2009).

65. Murtaugh, L. C., Stanger, B. Z., Kwan, K. M. & Melton, D. A. Notch signaling controls multiple steps of pancreatic differentiation. PNAS 100, 14920–14925 (2003).

66. Heng, X., Guo, Q., Leung, A. W. & Li, J. Y. Analogous mechanism regulating formation of neocortical basal radial glia and cerebellar Bergmann glia. eLife 6, e23253 (2017).

67. Hagan, N. & Zervas, M. Wnt1 expression temporally allocates upper rhombic lip progenitors and defines their terminal cell fate in the cerebellum. Molecular and Cellular Neuroscience 49, 217–229 (2012).

68. Morales, D. & Hatten, M. E. Molecular Markers of Neuronal Progenitors in the Embryonic Cerebellar Anlage. Journal of Neuroscience 26, 12226–12236 (2006).

69. Pierce, E. T. Histogenesis of the deep cerebellar nuclei in the mouse: an autoradiographic study. Brain Res 95, 503–518 (1975).

70. Green, M. J. et al. Independently specified Atoh1 domains define novel developmental compartments in rhombomere 1. Development 141, 389–398 (2014).

71. Imayoshi, I. & Kageyama, R. bHLH Factors in Self-Renewal, Multipotency, and Fate Choice of Neural Progenitor Cells. Neuron 82, 9–23 (2014).

72. Imayoshi, I. et al. Oscillatory Control of Factors Determining Multipotency and Fate in Mouse Neural Progenitors. Science 342, 1203–1208 (2013).

73. Yeung, J. et al. Wls Provides a New Compartmental View of the Rhombic Lip in Mouse Cerebellar Development. J. Neurosci. 34, 12527–12537 (2014).

74. Machold, R. P., Kittell, D. J. & Fishell, G. J. Antagonism between Notch and bone morphogenetic protein receptor signaling regulates neurogenesis in the cerebellar rhombic lip. Neural Development 2, 5 (2007).

75. Leung, A. W. & Li, J. Y. H. The Molecular Pathway Regulating Bergmann Glia and Folia Generation in the Cerebellum. Cerebellum 17, 42–48 (2018).

76. Li, K., Leung, A. W., Guo, Q., Yang, W. & Li, J. Y. H. Shp2-Dependent ERK Signaling Is Essential for Induction of Bergmann Glia and Foliation of the Cerebellum. J. Neurosci. 34, 922–931 (2014).

77. Altman, J. & Bayer, S. A. Development of the Cerebellar System: In Relation to Its Evolution, Structure, and Functions. (CRC-Press, 1997).

78. Mecklenburg, N. et al. Growth and differentiation factor 10 (Gdf10) is involved in Bergmann glial cell development under Shh regulation. Glia 62, 1713–1723 (2014).

79. Yamada, K. & Watanabe, M. Cytodifferentiation of bergmann glia and its relationship with purkinje cells. Anato Sci Int 77, 94–108 (2002).

80. Aldinger, K. A. & Doherty, D. The genetics of cerebellar malformations. Seminars in Fetal and Neonatal Medicine 21, 321–332 (2016).

81. Kopinke, D. et al. Ongoing Notch signaling maintains phenotypic fidelity in the adult exocrine pancreas. Developmental Biology 362, 57–64 (2012).

82. Zervas, M., Millet, S., Ahn, S. & Joyner, A. L. Cell Behaviors and Genetic Lineages of the Mesencephalon and Rhombomere 1. Neuron 43, 345–357 (2004).

83. Schmidt, U., Weigert, M., Broaddus, C. & Myers, G. Cell Detection with Star-convex Polygons. arXiv:1806.03535 [cs] 11071, 265–273 (2018).

84. Pécot, T., Cuitiño, M. C., Johnson, R. H., Timmers, C. & Leone, G. Deep learning tools and modeling to estimate the temporal expression of cell cycle proteins from 2D still images. bioRxiv 2021.03.01.433386 (2021) doi:10.1101/2021.03.01.433386.

85. Stuart, T. et al. Comprehensive Integration of Single-Cell Data. Cell 177, 1888–1902.e21 (2019).

86. Haghverdi, L., Lun, A. T. L., Morgan, M. D. & Marioni, J. C. Batch effects in single-cell RNA-sequencing data are corrected by matching mutual nearest neighbors. Nature Biotechnology 36, 421–427 (2018).

87. Korsunsky, I., Nathan, A., Millard, N. & Raychaudhuri, S. Presto scales Wilcoxon and auROC analyses to millions of observations. bioRxiv 653253 (2019) doi:10.1101/653253.

88. Yu, G., Wang, L.-G., Han, Y. & He, Q.-Y. clusterProfiler: an R Package for Comparing Biological Themes Among Gene Clusters. OMICS: A Journal of Integrative Biology 16, 284–287 (2012).

89. Granja, J. M. et al. ArchR is a scalable software package for integrative single-cell chromatin accessibility analysis. Nature Genetics 53, 403–411 (2021).

90. Pohl, A. & Beato, M. bwtool: a tool for bigWig files. Bioinformatics 30, 1618–1619 (2014).

91. Gel, B. et al. regioneR: an R/Bioconductor package for the association analysis of genomic regions based on permutation tests. Bioinformatics 32, 289–291 (2016).

92. van Heeringen, S. J. & Veenstra, G. J. C. GimmeMotifs: a de novo motif prediction pipeline for ChIP-sequencing experiments. Bioinformatics 27, 270–271 (2011).

